# Death-seq identifies regulators of cell death and senolytic therapies

**DOI:** 10.1101/2022.04.01.486768

**Authors:** Alex Colville, Jie-Yu Liu, Samantha Thomas, Heather D. Ishak, Ronghao Zhou, Julian D.D. Klein, David W. Morgens, Armon Goshayeshi, Jayesh S. Salvi, David Yao, Kaitlyn Spees, Michael C. Bassik, Thomas A. Rando

## Abstract

Selectively ablating senescent cells (“senolysis”) is an evolving therapeutic approach for age-related diseases. Current senolytics are limited to local administration by potency and side effects. While genetic screens could identify senolytics, current screens are underpowered for identifying genes that regulate cell death due to limitations in screen methodology. Here, we establish Death-seq, a positive selection CRISPR screen optimized to identify enhancers and mechanisms of cell death. Our screens identified synergistic enhancers of cell death induced by the known senolytic ABT-263, a BH3 mimetic. SMAC mimetics, enhancers in our screens, synergize with ABT-199, another BH3 mimetic that is not senolytic alone, clearing senescent cells in models of age-related disease while sparing human platelets, avoiding the thrombocytopenia associated with ABT-263. Death-seq enables the systematic screening of cell death pathways to uncover molecular mechanisms of regulated cell death subroutines and identify drug targets for diverse pathological states such as senescence, cancer, and neurodegeneration.

## INTRODUCTION

Cellular senescence is a cell state triggered by various stresses and characterized by prolonged irreversible cell-cycle arrest with secretory features (Collado et al., 2007; Di Micco et al., 2021; Gorgoulis et al., 2019). Elimination of senescent cells extends median lifespan of mice (Baker et al., 2016) and is being developed as a means to treat many age-related diseases (He et al., 2017) and to restore tissue homeostasis in aging (Xu et al., 2018; Baar et al., 2017). Senolytics, a class of drugs that selectively target senescent cells for death, hold promise for achieving those goals. However, the systemic toxicity and limited potency of current senolytic targets limit their use to local administration and compromise their clinical application (van Deursen, 2019).

Pooled genetic screens carry potential to identify improved senolytic targets. A recent RNAi approach has been used to identify a new senolytic target (Johmura et al., 2021), but this study failed to identify more than one senolytic target and any hits in classical cell death pathways in part because of the limitations of negative selection growth screens and RNAi interference screens. Negative selection growth screens on cells that rarely divide are even more challenging than on transformed cancer cells as the signal-to-noise ratios are decreased due to the lack of population doubling throughout the screen, which typically last 10-14 days in the case of negative selection growth screens. In contrast, an Annexin V-based enrichment screen outside of the senescence field pioneered the concept of a positive selection screen for dying cells (Arroyo et al., 2016). However, performing screens by selection of Annexin V+ cells requires either magnetic columns, which are not suitable for large cells such as senescent cells, or fluorescence-activated cell sorting, which suffers from limited throughput for genome-scale screens with time sensitive readouts like cell death.

To overcome these challenges and to increase the likelihood of uncovering targets in cell death pathways, we developed a method we termed “Death-seq” to positively select dying cells in response to genetic or pharmacologic interventions. Recent viability screens using primary neurons (Kramer et al., 2018) and muscle cells (Lek et al., 2020) highlight a growing opportunity to conduct genome-scale screens in more physiologically relevant cell culture models that are not as amenable to scale-up as immortalized cell lines. Yet applications of genetic screening to these models, such as in neurological diseases, remain scarce due in part to methodology limitations (So et al., 2019). Death-seq amplifies signal and reduces the noise compared to traditional dropout screens by design to further enable screening in more physiologically relevant models. We then implemented this approach with a genome-wide CRISPR screen to identify targets to enhance senolytic therapy.

We started with the senolytic therapy ABT-263, an inhibitor of the anti-apoptotic proteins BCL-2, BCL-xL (BCL2L1), and BCL-w (BCL2L2). Senescent cells are resistant to apoptosis which is thought to be mediated in part by the dysregulation of BCL-2 family (Yosef et al., 2016; Di Micco et al., 2021). The anti-apoptotic BCL-2 family proteins neutralize BAX and BAK to prevent their oligomerization and inhibit the BH3-only pro-apoptotic proteins that can activate BAX and BAK, thus inhibiting mitochondrial outer membrane permeabilization (MOMP). These interactions that regulate MOMP and apoptosis are dictated by the BCL-2 homology 3 (BH3) domain, and small molecule BH3 mimetics, such as ABT-263, were developed to enhance apoptosis. As a result of senescent cells dysregulation of the BCL-2 family, ABT-263 has been used as a senolytic both in vitro and in vivo (Zhu et al., 2016; Chang et al., 2016). Second mitochondria-derived activator of caspases (SMAC, also referred to as DIABLO) is released upon MOMP and inactivates IAPs, the inhibitor of apoptosis proteins, to promote apoptosis. The IAP protein family is known to regulate cell survival and three of the eight IAP family members have documented anti-apoptotic roles: X-chromosome-linked IAP (XIAP), cellular IAP 1 and 2 (cIAP1 and cIAP2) (Morrish et al., 2020). They interfere with apoptosis signaling via direct and indirect inhibition of caspases 3, 7, and 9. As a natural antagonist of IAPs, SMAC binds to IAPs to prevent their binding to caspases, leading to the activation of the downstream caspase cascade. Like BH3 mimetics, small molecule SMAC mimetics were developed to selectively enhance the death of cancer cells. Our screens systematically identify components of the intrinsic apoptosis pathway, ascertain uncharacterized regulators of senolysis, and uncover a synergy between BH3 mimetics and SMAC mimetics in senescent cell elimination with therapeutic potential.

## RESULTS

### Genome-wide senolytic CRISPR screen using Death-seq

We conducted a genome-wide CRISPR knockout (KO) screen in IMR-90 normal human lung fibroblasts and induced them to senescence by doxorubicin (Doxo-SEN) to discover knockouts that would exhibit either increased or decreased sensitivity to senolysis by ABT-263 (Figure 1A). After incubating the cells with ABT-263, also known as navitoclax, for 24 hours and collecting the dying floating and attached live cells, the sgRNA distributions of each population were determined by sequencing (“Death-seq”). The casTLE algorithm (Morgens et al., 2016) was used to estimate the effect and P value of each gene knockout. Death-seq enables the separate processing of the dying and live cells from the same plates, allowing the direct comparison of sgRNA distributions in each as opposed to comparing to a baseline frequency before the experiment or a DMSO-treated control arm, thereby amplifying signal and cutting tissue culture reagents in half.

**Figure 1.**
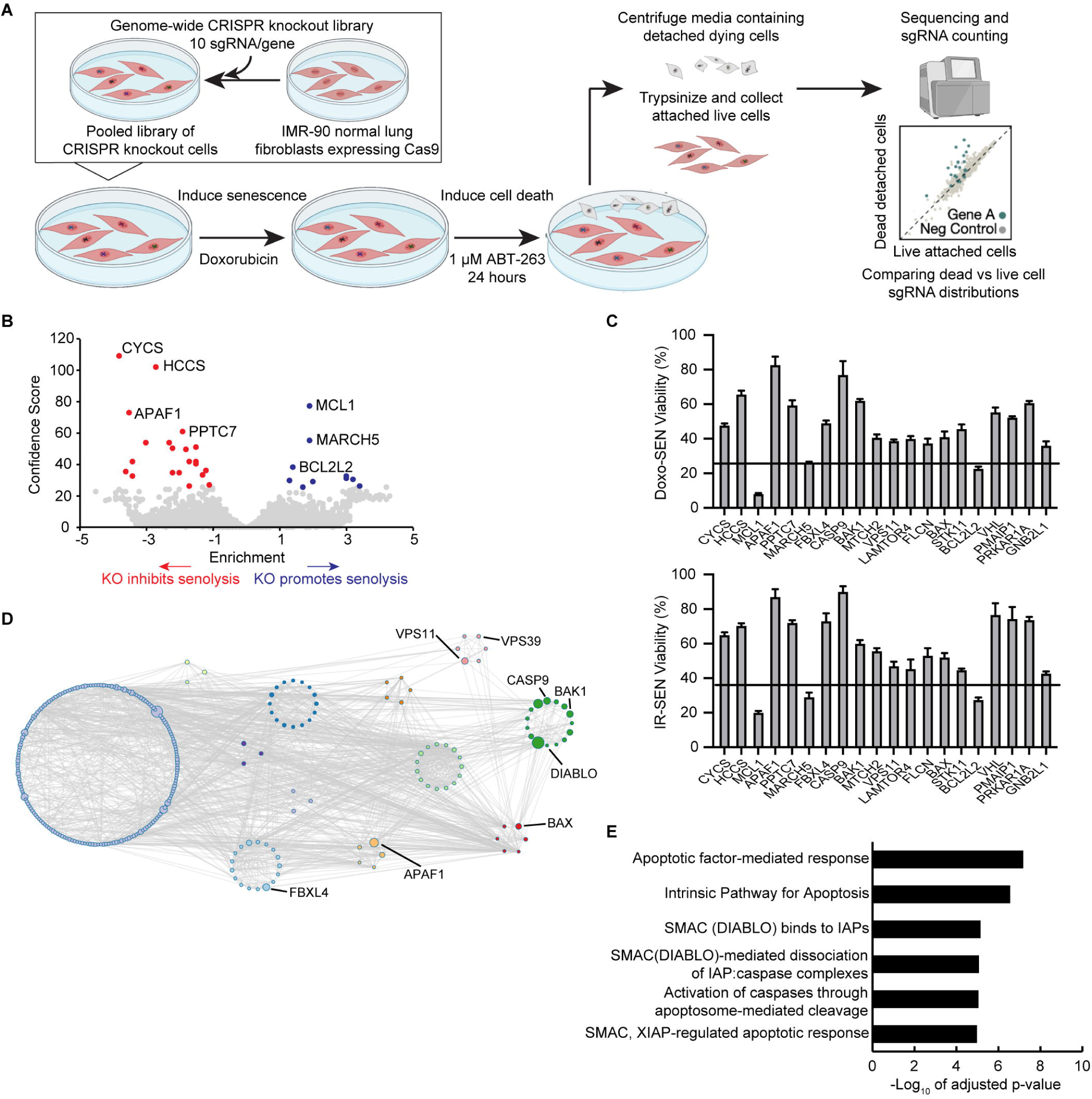
A genome-wide CRISPR screen for modifiers of cell death, Death-seq, identifies modifiers of senescent cell death. (A) Schematic of Death-seq screening method for genetic modifiers of cell death. The screen was performed in duplicate. (B) Volcano plot of the effects and confidence scores of all the genes in the genome-wide CRISPR screen in doxorubicin-induced senescent IMR-90s treated with 1 μM ABT-263 for 24 h. Effects and casTLE scores are calculated by casTLE. Labelled are the 31 genes passing 10% FDR for conferring resistance to (red) or promoting (blue) cell death by ABT-263 when knocked out. (C) Validation of the top 20 genome-wide screen hits using individual-well sgRNA knockouts of the indicated genes compared to control gRNA viability (represented by the line) after treatment with 1 μM ABT-263 for 3 d in either doxorubicin (Doxo)-induced (top) or irradiation (IR)-induced (bottom) senescent IMR-90s. Data are representative of two independent experiments performed in triplicate and are presented as mean□± □s.e.m. (D) Visualization of the genetic interaction network of the top 350 enriched genes in the genome-wide ABT-263 screen using Metascape. The size of the nodes corresponds to the confidence score of each gene in the screen. The Molecular Complex Detection (MCODE) algorithm was used to identify densely connected network components identified by circles of colored nodes. Select representative genes are highlighted. (E) The top Reactome categories enriched in the 64 genes that passed 30% FDR in the genome-wide ABT-263 screen. See also Figure S1 and the results and raw sequencing counts from screen in Tables S1 and S2 respectively.

Using a 30% false discovery rate (FDR) cutoff, the genome-wide screen identified 64 genetic modifiers which either inhibit or promote the senolytic effect of ABT-263 and 31 hits passing 10% FDR (Figure 1B; Tables S1 and S2). We validated the top 20 hits from the screen using sgRNA KO IMR-90s (Figure 1C). The top resistance hits including CYCS, HCCS, and APAF1 served as positive controls given their roles in the intrinsic apoptosis pathway. The fourth resistance hit, PPTC7, a mitochondrial phosphatase localized to the matrix, is poorly characterized (Niemi et al., 2019) with no previous connection to apoptosis and was also validated using shRNA knockdowns (Figure S1A). Consistent with recent studies (Li et al., 2020; Shahbandi et al., 2020; Han et al., 2017), MCL1, another BCL-2 family anti-apoptotic protein which is not targeted by ABT-263, mediated resistance to ABT-263. Unlike this recent work, combination treatment of the MCL1 inhibitor S63845 and ABT-263 had a stronger effect on proliferative cells (PROs) compared to senescent cells, as represented by Bliss score (Figures S1B and S1C). VHL, one of the top 20 hits, was further validated as a resistance hit given that it had an available small molecule inhibitor (Figure S1D). VHL encodes for the protein Von Hippel–Lindau tumor suppressor which functions as a E3 ubiquitin ligase and has been shown to regulate apoptosis (Li and Kim, 2011). Genetic interaction networks for the top 350 genes and pathway analysis for the 64 hits passing 30% FDR revealed a strong enrichment for pathways related to the intrinsic apoptosis pathway, mTOR signaling, and, unexpectedly, SMAC-IAP-caspase interactions (Figures 1D and 1E; Figure S1E).

### Death-seq head-to-head versus negative selection viability screens

In order to validate the genome-wide screen findings, we constructed a custom lentiviral sgRNA library targeting 384 genes to test the top ∼350 genome-wide screen hits as well as select apoptosis and lysosomal related genes (Table S3). We tested Death-seq head-to-head against a traditional negative selection screen design, which used a DMSO-treated control arm, using both an apoptosis sublibrary targeting ∼3000 genes as well as the newly constructed custom sgRNA library (Figure 2A). Death-seq identified 3 more hits reaching a 10% FDR threshold in the apoptosis sublibrary (Figure 2B) and 49 more hits in the custom sublibrary (Figure 2C) compared to the traditional method while achieving higher correlations between technical replicates. We then plotted the Death-seq combo confidence scores for each gene against those generated with the traditional method to compare the level of signal in the output of each method using both the apoptosis sublibrary (Figure 2D) and the custom sublibrary (Figure 2E). With the new positive selection method comparing live cells directly against dying cells, noise is reduced and signal is amplified enabling higher confidence scores and more significant hit calling. Next, to see if the method could identify senolytic hits even in the absence of a small molecule treatment, we used Death-seq on the DMSO-treated plates infected with our custom sublibrary. To our surprise, given the low screen coverage used, the well-known senolytic target BCL2L1 had the strongest casTLE score (Figure 2F).

**Figure 2.**
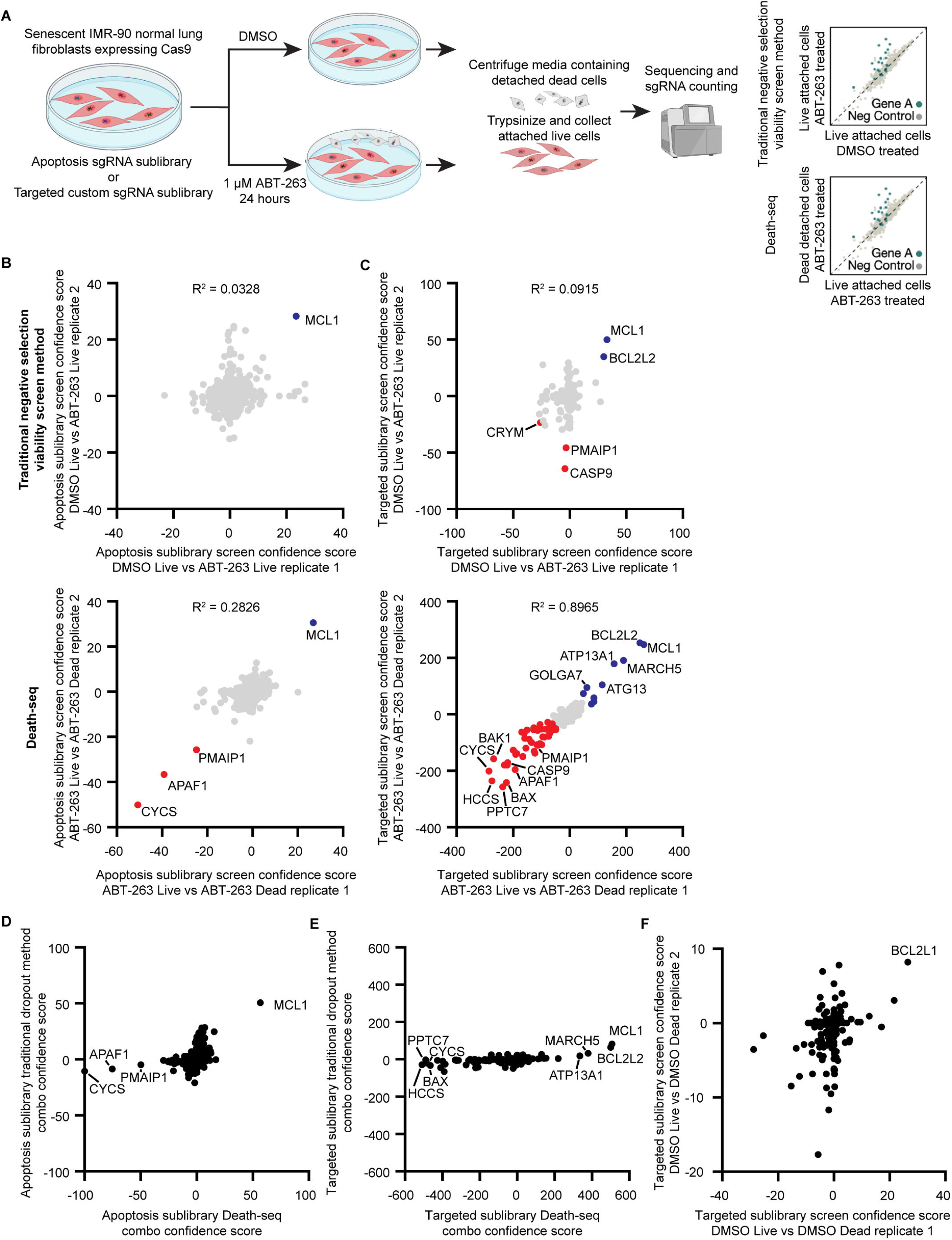
Head-to-head comparison of Death-seq against traditional negative selection viability screens. (A) Schematic of head-to-head comparisons of Death-seq screening method against traditional negative selection viability screens for genetic modifiers of cell death induced by ABT-263 treatment. The screens were performed in duplicate. (B and C) Correlation of casTLE confidence scores for all genes in the apoptosis (ACOC) (B) or targeted custom sublibrary (C) ABT-263 screen technical replicates comparing a traditional negative selection viability screen (top, DMSO-treated live cells vs. ABT-263-treated live cells) against Death-seq (bottom, ABT-263-treated live cells vs. ABT-263-treated dying cells). Labelled are the genes passing 10% FDR for conferring resistance to (red) or promoting (blue) cell death by ABT-263 when knocked out. R-squared values are from linear regression models. (D and E) Comparison of combo casTLE confidence scores generated using a traditional negative selection viability screen method (DMSO-treated live cells vs. ABT-263-treated live cells) vs. the Death-seq method (ABT-263-treated live cells vs. ABT-263-treated dying cells) for all genes in the apoptosis (ACOC) (D) or the targeted custom sublibrary (E). (F) Correlation of casTLE confidence scores for all genes in the targeted custom sublibrary DMSO screen technical replicates generated using the Death-seq method (DMSO-treated live cells vs. DMSO-treated dying cells). See also the results, raw sequencing counts, and composition of the targeted custom sublibrary from screens in Tables S1, S2, and S3 respectively.

### Targeted sublibrary senolytic Death-seq screens highlight importance of SMAC

Given the enrichment of pathways in the genome-wide screen related to the IAP-binding mitochondrial protein SMAC/DIABLO, we validated DIABLO as a resistance hit in sgRNA KO senescent IMR-90 cells (Figures 3A and S2A). To further validate the genome-wide ABT-263 screen and in hopes of finding a systemic senolytic therapy, we conducted parallel screens using ABT-263, the SMAC mimetic birinapant, and ABT-199 in Doxo-SENs as well as an additional ABT-263 screen in hydrogen peroxide-induced senescent cells (H_2_O_2_-SENs) (Figure 3B). Due to the well-documented on-target toxicity of ABT-263 to platelets limiting its use as a systemic therapy (Schoenwaelder et al., 2011), ABT-199, also known as venetoclax, was developed with enhanced BCL-2 specificity to avoid platelet toxicity (Souers et al., 2013). ABT-199 has been shown to not be senolytic in fibroblasts (Yosef et al., 2016). The Doxo-SEN ABT-263 custom sublibrary screen identified 80 hits passing a 10% FDR threshold (Figure 3C), including DIABLO and other members of the intrinsic apoptosis pathway, and the casTLE confidence scores of each gene correlated well with those of the genome-wide screen (Figure S2B). The H_2_O_2_-SEN and Doxo-SEN ABT-263 screens correlated strongly, potentially suggesting the two inducers yield similar senescence states (Figure S2C). The birinapant Doxo-SEN screen identified 14 hits which passed 10% FDR, 5 of which also passed 10% FDR in the ABT-263 screen (Figure 3D). Given their different mechanisms of action (MoAs), the birinapant and ABT-263 screens had a weak correlation. In support of the idea that these hits are combinatorial, BCL2L1 was a hit in the birinapant screen and DIABLO was a hit in the ABT-263 screen. The ABT-199 and ABT-263 screens had a strong correlation given their similar MoAs (Figure 3E). The ABT-199 screen identified 58 hits which passed 10% FDR including both DIABLO and XIAP.

**Figure 3.**
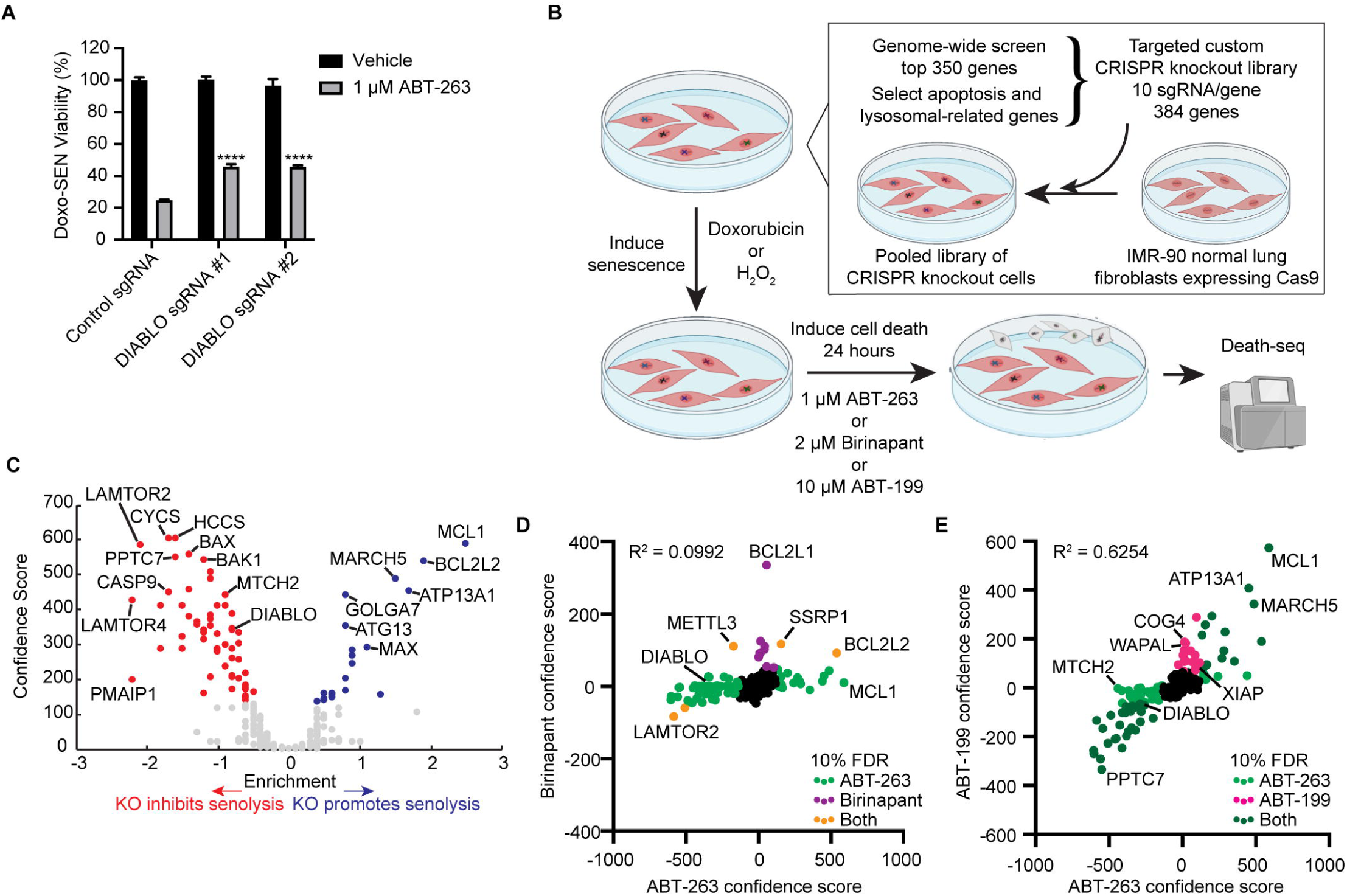
Targeted sublibrary senolytic Death-seq screens highlight importance of SMAC. (A) Validation of DIABLO/SMAC sgRNA knockouts compared to control gRNA viability after treatment with vehicle or 1 μM ABT-263 for 3 d in Doxo-induced senescent IMR-90s. Data are representative of two independent experiments performed in triplicate and are presented as mean□±□s.e.m. One-way ANOVA with Dunnett’s post-hoc test relative to control sgRNA-treated cells in the corresponding drug treatment, *P < 0.05, **P < 0.01, ***P < 0.001, ****P < 0.0001. (B) Schematic of targeted custom sublibrary screens. The targeted custom library was infected using lentivirus into Cas9-expressing IMR-90 normal lung fibroblasts. The IMR-90s expressing the targeted sublibrary were then induced to senesce with Doxo and treated with 1 μM ABT-263, 2 μM birinapant, or 10 μM ABT-199 for 24 h. A targeted custom sublibrary screen was also performed with H_2_O_2_-induced senescent IMR-90s and 1 μM ABT-263 treatment. Like the genome-wide screen, Death-seq was used to compare the dying cell populations directly with the live attached cells. The screens were performed in duplicate. (C) Volcano plot of the effects and confidence scores of all the genes in the targeted custom CRISPR screen in Doxo-induced senescent IMR-90 cells treated with 1 μM ABT-263 for 24 h. Labelled are the 80 genes passing 10% FDR for conferring resistance to (red) or promoting (blue) cell death by ABT-263 when knocked out. (D) Correlation of combo casTLE confidence scores of the ABT-263 and birinapant screens in Doxo-induced senescent IMR-90s for all genes in the targeted custom sublibrary. R-squared value is from a linear regression model. Labelled in light green, purple, and orange are hits passing 10% FDR only in ABT-263 screen, only in birinapant screen, or in both screens, respectively. (E) Correlation of combo casTLE confidence scores of the ABT-263 and ABT-199 screens in Doxo-induced senescent IMR-90s for all genes in the targeted custom sublibrary. R-squared value is from a linear regression model. Labelled in light green, pink, and dark green are hits passing 10% FDR only in ABT-263 screen, only in ABT-199 screen, or in both screens, respectively. See also Figure S2 and the results, raw sequencing counts, and composition of the targeted custom sublibrary from screens in Tables S1, S2, and S3 respectively.

To further visualize the results of the Doxo-SEN ABT-263 custom sublibrary screen, we created a subcellular localization map with selected hits passing the 10% FDR threshold (Figure S2D). Many hits involved mitochondrial biology as previously discussed, serving as a positive control for the ability of the Death-seq method to identify components of cell death subroutine pathways. MARCH5, an E3 ubiquitin ligase known to mediate the degradation of proteins that regulate mitochondrial fission and fusion (Yonashiro et al., 2006), was one of the mitochondrially located hits which promoted senolysis by ABT-263 and ABT-199 in agreement with recent work in cancer cells (Arai et al., 2020; Subramanian et al., 2016). Recently, additional evidence was published validating our screen findings and revealing the likely mechanism through which our resistance screen hit MTCH2 assists in senolysis by aiding MARCH5 in turnover of the MCL1:NOXA complex (Djajawi et al., 2020).

As highlighted in the pathway analysis of the genome-wide screen results (Figure S1E), we identified a signal for mTOR signaling with several hits involving lysosomal biology. Our sublibrary screens identified several members of the lysosomal membrane–anchored Rag-Ragulator complex, a regulator of cellular responses to nutrient availability and metabolism. These members include FLCN, all five LAMTOR subunits 1-5, RRAGA, and RRAGC as resistance hits to senolysis induced by ABT-263. Interestingly, a role of the Rag-Ragulator complex in cell death was not uncovered until a recently published CRISPR screen identified the mechanistic role of the complex in pyroptosis (Zheng et al., 2021). However, this work did not find a role for the complex in apoptosis. Our ABT-263 senescence screens also identified hits involving the lysosomal associated GATOR complex, known for amino acid sensing. These included synergistic hits within the GATOR1 subcomplex, including NRPL2 and NPRL3, and resistance hits in the GATOR2 subcomplex, including WDR24, WDR59, SEH1L, and MIOS. Furthermore, the screens identified resistance hits in the lysosomal associated tethering complexes HOPS and CORVET including VPS16, VPS18, VSP11, VPS41, and VPS39. The breadth and systematic nature of the identified modifiers of senolysis further validate our screening method.

### BH3 and SMAC mimetics synergize to induce senolysis

We next focused on the intriguing, recurring resistance hit SMAC/DIABLO. Consistent with our genetic screening results, we observed a synergistic inhibition of cell viability in Doxo-SEN IMR-90s using the combination of ABT-263 and SMAC mimetic birinapant as calculated by the Bliss independence model (Figures 4A and 4B) and no synergistic relationship in proliferative cells (Figure S3A). Likewise, the combination of ABT-263 with other SMAC mimetics SM-164 and GDC-0152 also produced this increased senolytic effect (Figures S3B and S3C). We also validated the hits DIABLO and XIAP from the ABT-199 screen with the combination of ABT-199 and birinapant showing an even stronger synergy on senescent cell viability as shown by Bliss score (Figures 4C and 4D) while still sparing proliferative cells (Figure S3D). This result was further validated using ABT-199 in combination with SM-164 or GDC-0152 (Figures S4A and S4B), as well as in IR-SENs with all three SMAC mimetics (Figures S4C-S4E).

**Figure 4.**
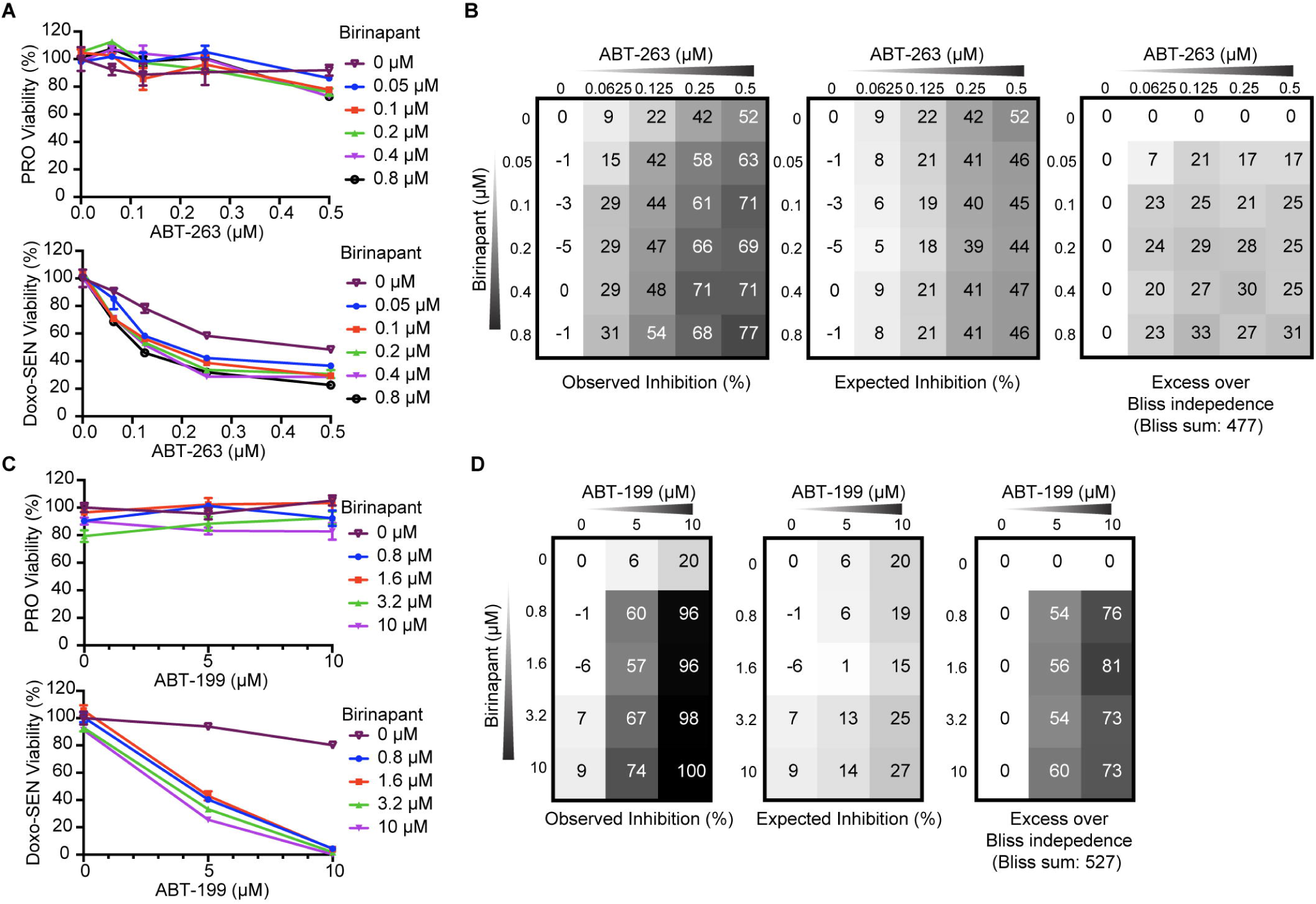
BH3 and SMAC mimetics synergize to induce selective death in senescent cells. (A) Proliferative (top) and Doxo-induced senescent (bottom) IMR-90s were treated with ABT-263 and birinapant at the indicated concentrations for 3 d before viability was assessed relative to no drug control. Data are representative of two independent experiments performed in triplicate and are presented as mean□± □s.e.m. (B) The percent expected inhibition is subtracted from the percent observed inhibition at each combination of drug doses in Doxo-induced senescent IMR-90s to calculate drug synergy represented by excess over Bliss independence. (C) Proliferative (top) and Doxo-induced senescent (bottom) IMR-90s were treated with ABT-199 and the SMAC mimetic birinapant at the indicated concentrations for 3 d before viability was assessed relative to no drug control. Data are representative of two independent experiments performed in triplicate and are presented as mean□± □s.e.m. (D) The percent expected inhibition is subtracted from the percent observed inhibition at each combination of drug doses in Doxo-induced senescent IMR-90s to calculate drug synergy represented by excess over Bliss independence. See also Figure S3 and S4.

Having validated the ability of ABT-199 and SMAC mimetics to synergize in senolysis, we next sought to orthogonally confirm cell death, mitochondrial outer membrane permeabilization, and the senolytic ability of the combination in different inducers of senescence and cell types. Using flow cytometry and Annexin V staining, we analyzed the extent of cell death in proliferative and senescent IMR-90s treated with vehicle, ABT-199/SMAC mimetic, or ABT-263, each with or without the pan-caspase inhibitor Q-VD-Oph (QVD). As with ABT-263, the percent of apoptotic cells in the ABT-199/SMAC mimetic group decreased with QVD treatment, suggesting that the combination acts through caspase-mediated apoptosis in senescent cells (Figure 5A). Mitochondrial membrane potential was significantly reduced in senescent cells treated with ABT-199 alone, ABT-199/SMAC mimetic, and ABT-263, but not in proliferative cells (Figure 5B). Caspase 3/7 activity was significantly increased in senescent cells treated with ABT-199/SMAC mimetic and ABT-263, but not in proliferative cells (Figure 5C). The combination of ABT-199 and SMAC mimetic was senolytic across different senescence inducers in IMR-90s including H_2_O_2_, bleomycin, palbociclib, and etoposide (Figure S5A). The combination was also senolytic and synergistic in senescent WI-38 normal lung fibroblasts (Figures 5D and S5B). We further tested the combination in senescent human retinal microvascular endothelial cells (HRMECs), a disease-relevant primary cell population recently utilized in a study which implicated the pathological role of senescent cells in diabetic retinopathy (Crespo-Garcia et al., 2021). ABT-199/birinapant was also senolytic in HRMECs (Figure S5C).

**Figure 5.**
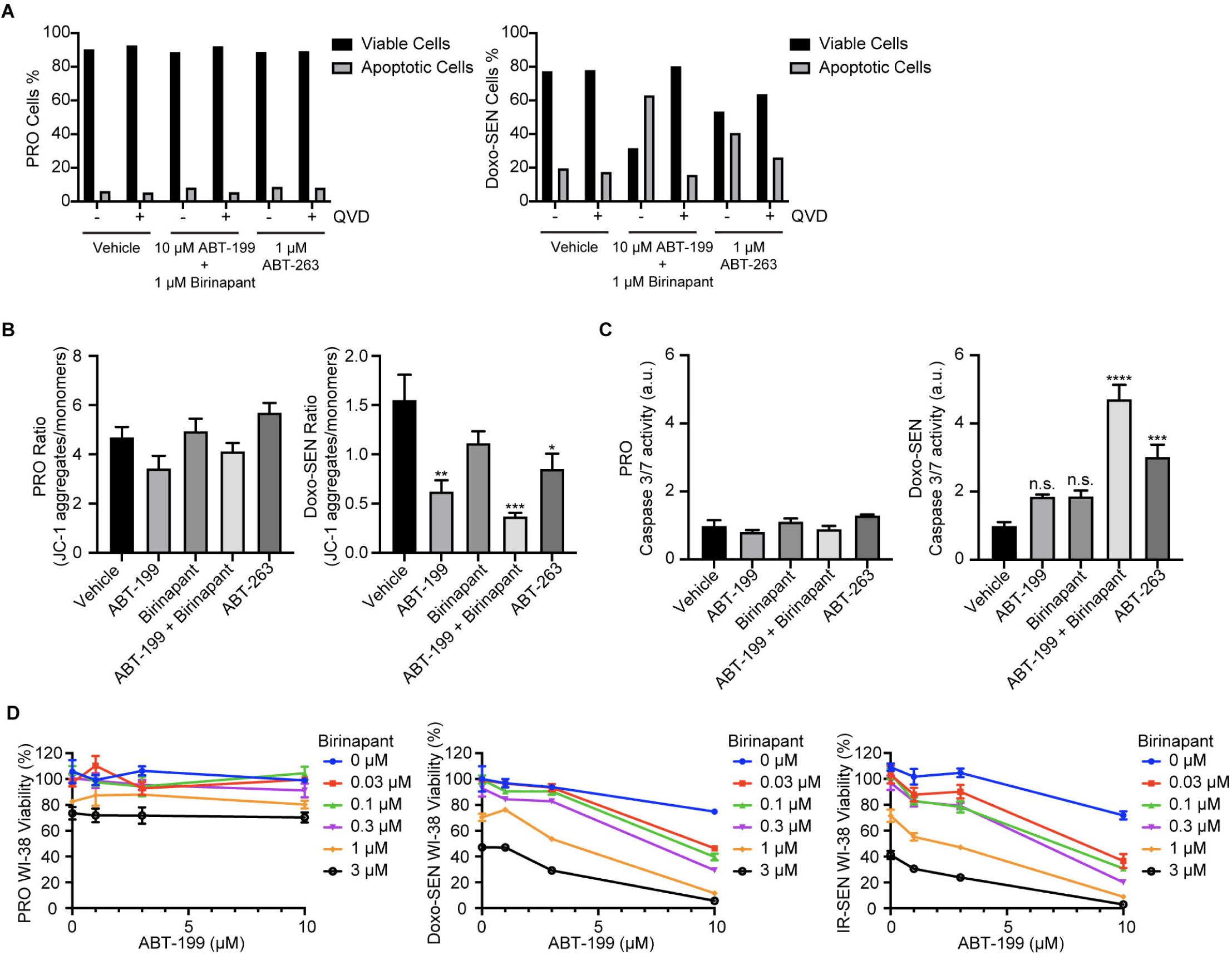
ABT-199 and SMAC mimetic combination is senolytic through apoptosis across different cell types. (A) Percentage of viable and apoptotic proliferative (left) and Doxo-induced senescent (right) IMR-90s 24 h after treatment with vehicle, 10 μM ABT-199 and 1 μM birinapant, or 1 μM ABT-263, all with and without 20 μM Q-VD-Oph (QVD). (B) Proliferative (left) and Doxo-induced senescent (right) IMR-90s were treated for 24 h with vehicle, 10 μM ABT-199, 1 μM birinapant, 10 μM ABT-199 and 1 μM birinapant, or 1 μM ABT-263. The treated cells were then stained and measured by flow cytometry to determine the ratio of JC-1 aggregates to JC-1 monomers. Data are representative of two independent experiments performed in triplicate and are presented as mean□± □s.e.m. One-way ANOVA with Dunnett’s post-hoc test, *P < 0.05, **P < 0.01, ***P < 0.001, ****P < 0.0001. (C) Normalized caspase 3/7 activity in proliferative (left) and Doxo-induced senescent (right) IMR-90s that were treated for 24 h with vehicle, 10 μM ABT-199, 1 μM birinapant, 10 μM ABT-199 and 1 μM birinapant, or 1 μM ABT-263. Data are representative of two independent experiments performed in triplicate and are presented as mean□± □s.e.m. One-way ANOVA with Dunnett’s post-hoc test, *P < 0.05, **P < 0.01, ***P < 0.001, ****P < 0.0001. (D) Proliferative (left), Doxo-induced senescent (middle), and IR-induced senescent (right) WI-38s were treated with ABT-199 and birinapant at the indicated concentrations for 3 d before viability was assessed relative to no drug control. See also Figure S5.

### ABT-199 and SMAC mimetic combination spares human platelets and reduces levels of senescent cell markers in vivo

Despite that BCL-2 is not required for platelet survival (Debrincat et al., 2015), the effect of simultaneous inhibition of BCL-2 and IAPs on platelets is still unclear. To determine if the combination of ABT-199 and birinapant is toxic to platelets, we cultured human platelets in either ABT-263, ABT-199, or the ABT-199/birinapant combination before assessing viability. ABT-263 significantly depleted platelets starting at 300 nM while the combination did not affect human platelet viability (Figures 6A, S6A, and S6B).

**Figure 6.**
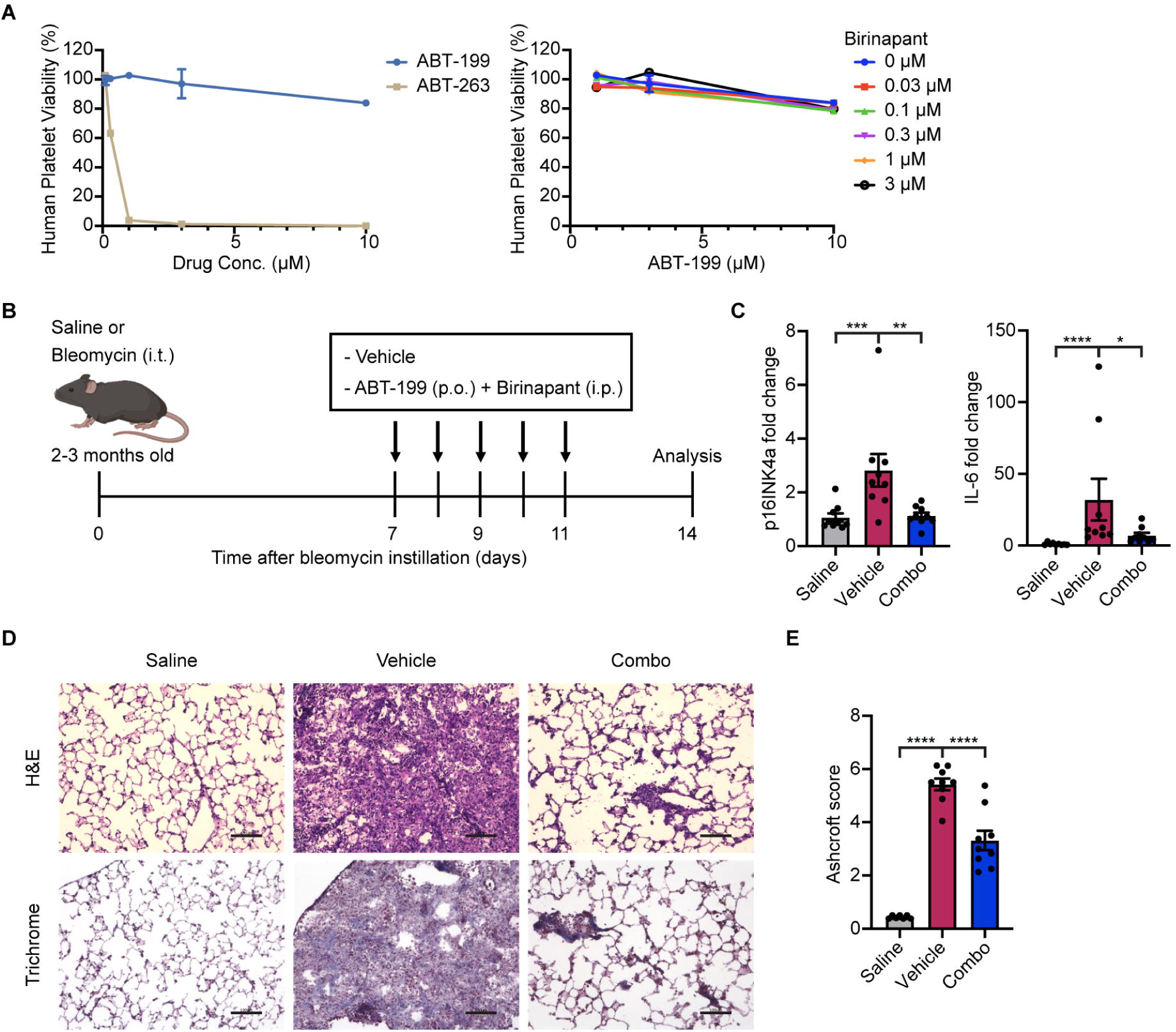
ABT-199 and SMAC mimetic combination spares human platelets and reduces levels of senescent cell markers in vivo. (A) Human platelets were treated with ABT-199 or ABT-263 (left) or ABT-199 in combination with birinapant at the indicated concentrations (right) for 3 d before viability was assessed relative to no drug control. Data are representative of two independent experiments performed in triplicate and are presented as mean□± □s.e.m. (B) Schematic of bleomycin-induced IPF mouse model experiment. (C) mRNA expression of *p16*^*INK4a*^ and *Il6* in murine lungs were quantified by qRT-PCR and normalized by TBP levels. Fold-increase was calculated with respect to the mRNA levels in saline-treated control mice (n = 8 in each group). Data are representative of two independent experiments performed in triplicate and are presented as mean□± □s.e.m. One-way ANOVA with Dunnett’s post-hoc test, *P < 0.05, **P < 0.01, ***P < 0.001, ****P < 0.0001. (D) Representative images of hematoxylin and eosin (top) and Masson’s trichrome (bottom) stained lung sections from saline control (left), vehicle-treated (middle), and ABT-199/birinapant-treated (right) mice. Scale bar: 100 μm. (E) Pulmonary fibrosis was quantified with modified Ashcroft scores (n = 7 control group, n = 9 vehicle and combo groups). Data are representative of two independent experiments performed in triplicate and are presented as mean□± □s.e.m. One-way ANOVA with Dunnett’s post-hoc test, *P < 0.05, **P < 0.01, ***P < 0.001, ****P < 0.0001. See also Figure S6.

Given the potency and selectivity of the ABT-199/birinapant combination against senescence in vitro and the absence of toxicity to human platelets, we wondered whether the combination could be used therapeutically in a systemic fashion. Recent evidence has suggested a causal role of senescent cells in the pathogenesis of idiopathic pulmonary fibrosis (IPF) (Schafer et al., 2017; Yao et al., 2020; Merkt et al., 2020; Liu and Liu, 2020). IPF is a deadly chronic lung disease characterized by progressive lung scarring with a median survival of less than three years and aging as a major risk factor (Schneider et al., 2021; Martinez et al., 2017; Barnes, 2013). Consistently, studies have demonstrated an accumulation of senescent cells in human IPF lungs (Schafer et al., 2017; Martinez et al., 2017; Álvarez et al., 2017; Adams et al., 2020). Moreover, the importance of human genetic variants in telomerase in both sporadic and familial disease suggests the pathological contribution of cellular senescence (Moore et al., 2019; Armanios et al., 2007; Povedano et al., 2015). Removal of senescent cells in experimental lung fibrosis models alleviated pulmonary fibrosis and improved lung function (Schafer et al., 2017; Wiley et al., 2019; Pan et al., 2017). We therefore tested the ability of the systemic combination to clear senescent cells in the bleomycin-injury IPF model, in which bleomycin induces senescence in various lung cells including alveolar epithelial cells and myofibroblasts (Schafer et al., 2017; Aoshiba et al., 2003; Hecker et al., 2014). Mice were treated with a single dose of bleomycin through intratracheal administration and subsequently treated with vehicle or the ABT-199/birinapant combination (Figure 6B). The levels of senescent cell markers including *p16*^*INK4a*^ and *Il6* increased 14 days after bleomycin injury and were significantly decreased by the combination treatment (Figure 6C). Strikingly, treatment with the combination significantly reduced the severity of pulmonary fibrosis assessed by histological staining and Ashcroft histopathology scoring (Figures 6D and 6E).

Next, we tested the combination in an independent disease model. Senescence is associated with metabolic disease and obesity-induced disorders including nonalcoholic steatohepatitis (NASH) (Palmer et al., 2019; Papatheodoridi et al., 2020). Targeting senescent cells has been shown to mitigate liver damage in mice receiving high-fat diet (Omori, et al., 2020; Johmura et al., 2021; Ogrodnik et al., 2017; Amor et al., 2020). In a mouse model of NASH induced by feeding of a choline-deficient, high-fat diet (CDA-HFD), mice were treated with vehicle, the ABT-199/birinapant combination, or ABT-263 after four weeks on a CDA-HFD (Figure S6C). Supporting the induction of senescence in the NASH model, levels of liver *p16*^*INK4a*^ mRNA and serum alanine aminotransferase (ALT) were significantly increased by the diet in the vehicle CDA-HFD group compared to normal chow controls (Figure S6D and S6E). We noted a reduction in liver *p16*^*INK4a*^ levels by the combination (Figure S6D), as well as a lower level of serum alanine aminotransferase (ALT), suggesting that liver function was improved (Figure S6E). Overall, this systemic senolytic combination treatment resulted in the reduction of senescent cell markers consistent with their clearance and partially ameliorated some aspects of age-related pathologies.

## DISCUSSION

Here, we present a screening method to systematically assess modifiers of cell death. We discovered a number of robust modifiers of ABT-263 senolytic therapy and demonstrated the ability to identify genetic modifiers of death, even with drugs that do not alone induce significant amounts of cell death such as ABT-199 and birinapant. Using the new screening method, we were able to generate a high-quality catalog of genes that served as modifiers of senescent cell death. These data sets of modifiers of senolysis will serve as an important resource for clinicians and experts in the fields of cell death, BH3 mimetics, SMAC mimetics, mitochondrial biology, and cellular senescence. The BCL-2 family of proteins has many intra-family interactions and a high degree of complexity that makes it difficult to dissect and predict cell fate with standard molecular biology techniques (Montero and Letai, 2018). Our results will help provide a resource and technique to further the molecular understanding of BCL-2 inhibitors, especially in the context of cellular senescence where there is a dearth of work relative to the cancer field. The data sets not only contain a number of well characterized members of the apoptosis pathway, but also less well characterized genes in apoptosis pathways such as HCCS which encodes for the enzyme holocytochrome c-type synthase. Many hits intriguingly have no previous connection to apoptosis, cell death, or senescence. An example of one of these hits is PPTC7, a strong resistance hit from both our ABT-263 and ABT-199 senolysis screens. PPTC7 encodes for a poorly characterized PP2C phosphatase localized to the mitochondrial matrix. Published work has revealed evidence for the role of PPTC7 in mitochondrial metabolism (Niemi et al., 2019). However, to our knowledge, no evidence exists to suggest a role of PPTC7 in cell death. Two screen hits which promoted senolysis, GOLGA7 and ATP13A1, are two more examples of surprising hits, given the lack of documented connection to senolysis or apoptosis, with GOLGA7 recently found to be necessary for an unconventional, non-apoptotic form of cell death (Ko et al., 2019). Our ABT-263 and ABT-199 screen resistance hits also pointed to the role of the Rag-Ragulator complex, the GATOR complex, the HOPS complex, and the CORVET complex in cell death. Future work is required to help tease apart the mysteries revealed by Death-seq, including the role of PPTC7, GOLGA7, ATP13A1, and the lysosomal associated complexes in senolysis by BH3 mimetics, as well as the many other hits not currently associated with cell death or senescence.

We validated the unexpected finding from our screens that the BH3 mimetics ABT-263 and ABT-199 act in a synergistic fashion with SMAC mimetics to induce senolysis. Preliminary in vitro studies in the cancer field hint at the potential of a synergy between BCL-2 inhibition and SMAC mimetics to improve efficacy and reduce toxicity (Perimenis et al., 2016; Chen et al., 2012). This work suggested that the dysregulation of the IAP proteins in cancer cells may hinder the death of cancer cells. Just like cancer cells, senescent cells are regarded as resistant to cell death due to their general dysregulation of anti-apoptotic proteins (Di Micco et al., 2020). However, the exact mechanisms of this anti-apoptotic phenotype remain to be investigated. Many questions remain to be answered regarding the molecular mechanisms of the synergy between ABT-199 and SMAC mimetics in senolysis and whether this relates to the general phenomenon of senescent cells resistance to death.

Senolytics have shown promise as a strategy to address age-related disease, but because of toxicities most senolytics have been limited to local administration instead of used to treat the body systemically. Given the world’s dramatically increasing elderly population, it is important to develop strategies to extend and restore healthspan (Kennedy et al., 2014; Olshansky 2018; Scott et al., 2021). The COVID-19 pandemic has demonstrated the enhanced vulnerability of the elderly and those with age-related disease to hospitalization, death, and other adverse events as a result of infections, in this case with severe acute respiratory syndrome coronavirus 2 (SARS-CoV-2). Senescent cells have been highlighted as a potential reason for this enhanced vulnerability of the elderly (Nehme et al., 2020) with a recent study demonstrating that senescent cells have an amplified inflammatory response to SARS-CoV-2 and that in old mice infected with SARS-CoV-2 related virus, senolytics reduce mortality and increase antiviral antibodies (Camell et al., 2021). A systemic senolytic treatment with minimal side effects could be used in the future to protect the elderly and those with age-related disease from SARS-CoV-2 and future pandemics. Likewise, a tolerable systemic senolytic treatment will be critical for clinical trials in other diseases that cannot be treated with local administration such as IPF, COPD, atherosclerosis, and neurodegeneration (van Deursen 2019). With this goal in mind, we find that the combination of ABT-199 and birinapant is not toxic to human platelets and reduces levels of senescent cell markers in mouse models of IPF and NASH establishing therapeutic potential. ABT-199 is FDA approved and birinapant has completed several Phase 2 clinical trials which could accelerate the clinical use of this combination. At least eight SMAC mimetics have shown to be well tolerated with reasonable safety profiles in human trials (Morrish et al., 2020). The reasons for which SMAC mimetics are inherently non-toxic to healthy tissue are still being investigated but are likely due to their targeted nature. The drug class has failed so far to advance to approval due to limited clinical activity as a single-agent in oncology.

Death-seq, through its comparison of guide composition in dying cells directly to live cells, allows more efficient and effective genome-wide screens and unlocks the potential for new findings in the senescence field and in a wide variety of fields outside of senescence. Due to the limitations of conventional negative selection screens, CRISPR screens have yet to be used to identify senolytic targets. Considering the pleiotropic phenotypes of senescent cells, other groups have stated the need to shift from traditional reductionistic scientific methods to new systematic high-throughput methods to study senescence (Gorgoulis et al., 2019). Similar to the cancer field, systematic efforts are needed to untangle the anti-apoptotic defense mechanisms of senescent cells. In addition to enabling the study of senescent cells, Death-seq can be of benefit to many other areas of biological and pathological study. The simplicity of the new screening method and its ability to increase signal and decrease noise will enable the identification of enhancers and mechanisms of cell death subroutines in other fields where it was previously not possible with currently available techniques. For instance, our method is well suited to contribute to work seeking to identify genetic modifiers of excess cell death in neurodegeneration and cardiovascular disease as well as mechanisms of drug resistance and synthetic lethality in cancer. Additionally, our method will improve the ability to screen in more physiologically relevant culture systems such as 3D organoid systems (Han et al., 2020), which are harder to scale-up and/or more resource intensive compared to immortalized cell lines.

### Limitations of the study

Although we provide evidence that combinations of BH3 and SMAC mimetics are senolytic across a variety of cell types and inducers of senescence, senescence is a heterogeneous state and as such we expect that the results from our screen will be to some extent specific to the cell states used in screening. As understanding of which senescent cell types play pathogenic versus beneficial or negligible roles in age-related disease and aging continues to improve, we expect that Death-seq will be utilized in additional cell types based on their disease-specific pathogenic relevance. Just like the Cancer Dependency Project has profiled hundreds of cancer cell models to identify genetic vulnerabilities for therapeutic development (Tsherniak et al., 2017), we anticipate that Death-seq will enable a similar project in the field of senescence across different models and inducers of senescence. Additionally, given the nascent tools which exist to study in vivo senescence (Roy et al., 2020), our mouse models of senescence are limited by the number of robust in vivo biomarkers available to confirm senolytic efficacy.

## ACKNOWLEDGMENTS

We thank S. Dixon for the helpful discussions and insights. We thank C. Cain, B. Carter, and the Palo Alto VA Flow Cytometry Core for assistance with flow cytometry experiments. A.C. thanks J. Pluvinage, R. Kamber, M. Schultz, Y. Lu, and M. Bonkowski (deceased) for scientific mentorship. We thank Y. Yu and the entire Rando and Bassik labs for laboratory assistance and discussion. Some illustrations were created with BioRender.com. We acknowledge our funding sources: the National Science Foundation Graduate Research Fellowship DGE-1656518 (A.C.), the Glenn Foundation for Medical Research (T.A.R.), the National Institutes of Health grant P01AG036695 (T.A.R.), the National Institutes of Health grant R01AR073248 (T.A.R.), and the Departments of Veterans Affairs BLR&D Merit Review I01BX002324 (T.A.R.).

## AUTHOR CONTRIBUTIONS

Conceptualization: A.C., J.-Y.L., J.D.D.K., D.W.M., M.C.B., T.A.R.

Methodology: A.C., J.-Y.L., S.T., H.D.I., J.D.D.K., D.W.M.

Investigation: A.C., J.-Y.L., S.T., H.D.I., J.D.D.K., D.W.M., R.Z., J.S.S., A.G., D.Y., K.S.

Visualization: A.C., J.-Y.L.

Funding acquisition: T.A.R., M.C.B., A.C.

Project administration: A.C., J.-Y.L., M.C.B., T.A.R.

Writing – original draft: A.C., J.-Y.L., T.A.R.

Writing – review & editing: A.C., J.-Y.L., T.A.R.

## STAR METHODS

### RESOURCE AVAILABILITY

#### Lead contact

Further information and requests for resources and reagents should be directed to and will be fulfilled by the lead contact, Thomas Rando.

#### Materials availability

This study did not generate new unique reagents.

#### Data and code availability

All screening data were analyzed with the casTLE scripts version 1.0 that are available at https://github.com/elifesciences-publications/dmorgens-castle. Flow cytometry data was analyzed using FlowJo 10 software. Statistics were calculated using GraphPad Prism 9 software. Any additional information required to reanalyze the data reported in this paper is available from the lead contact upon request.

### EXPERIMENTAL MODEL AND SUBJECT DETAILS

#### Cell culture

Media was changed every two to four days and cells were cultured in tissue culture plates in a humidified 37ºC incubator set at 5% CO_2_ and 5% O_2_. IMR-90s and WI-38s were cultured in Dulbecco’s Modified Eagle Medium (Corning, 10013CV) supplemented with 10% heat-inactivated FBS (Omega Scientific FB-02) and 1% pen-strep (Omega Scientific PS-20). HRMECs were cultured in Endothelial Cell Growth Medium MV2 kit (PromoCell, C-22022). Cells were periodically tested to ensure absence of mycoplasma. To induce cellular senescence by doxorubicin, cells were treated with doxorubicin at 250 nM for the first 24 hours then 83.3 nM for the subsequent 48 hours before washing out drug and culturing for another 4 days before experimentation. To induce cellular senescence by irradiation, cells were exposed to 10 Gy X-rays in an X-RAD 320 before changing the media and waiting 10 days before experimentation. To induce cellular senescence by H_2_O_2_, cells were treated with 200 μM H_2_O_2_ for 2 hours every other day for three total treatments and experimentation began 7 days after the first treatment. To induce cellular senescence with bleomycin, cells were treated with 40 μg/mL bleomycin for 2 hours and experimentation began 7 days after initial treatment. To induce cellular senescence with palbociclib, cells were treated with 10 μM palbociclib for 7 days before experimentation. To induce cellular senescence with etoposide, cells were treated with 50 μM etoposide for 48 hours and experimentation began 7 days after initial treatment.

#### Human platelet isolation and viability assay

Human platelets were isolated and cultured as described (He et al., 2020). In brief, human platelet-rich plasma (PRP) was purchased within 3 days after harvest. The PRP was transferred into a 50 mL conical tube with 5 mL acid citrate buffer (Santa Cruz Biotechnology, sc-214744). This mixture was centrifuged at 250g for 20 minutes 5 at room temperature without the centrifuge brake. PRP was collected into a 15 mL polypropylene conical tube and prostaglandin E1 (PGE1) (Santa Cruz Biotechnology, sc-201223A) and apyrase (Sigma-Aldrich, A6237) were added to final concentrations of 1 μM and 0.2 units/mL, respectively, to prevent clotting. After gently mixing the solution, platelets were pelleted by centrifugation at 1200g for 10 minutes without the centrifuge brake and the pelleted platelets were gently washed without disrupting the pellet in 2 mL HEPES Tyrode’s buffer (Boston BioProducts, PY-921WB) containing 1 μM PGE1 and 0.2 units/mL apyrase. After washing, pellets were slowly resuspended in 10 mL HEPES Tyrode’s buffer containing 1 μM PGE1, 0.2 units/mL apyrase, and 10% FBS. For the viability assays, platelet number was adjusted to 2 × 10^8^/mL in HEPES Tyrode’s buffer containing 1 μM PGE1, 0.2 units/mL apyrase, and 10% FBS. Each treatment was conducted in 2 mL of platelet suspension in 15 mL polypropylene conical tubes on a rotating platform at room temperature and the viability of platelets was measured after treatment for indicated time points by the previously described XTT assay. Extra precautions such as room temperature storage/incubation, centrifuging without a deceleration brake, gently pipetting and 20 mixing, and rotating platelets during incubation were taken to avoid platelet activation, aggregation, and spontaneous apoptosis.

#### Animals

Animal procedures were approved by the Administrative Panel on Laboratory Animal Care of the VA Palo Alto Health Care System. C57BL6 male mice were housed in specific pathogen-free conditions in barrier protected rooms under a twelve-hour light-dark cycle and were fed ad libitum.

#### In vivo treatment

ABT-263 and ABT-199 was prepared in 10% ethanol, 30% polyethylene glycol 400, and 60% Phosal 50 PG, followed by sonication at room temperature for 2 hours. Birinapant was prepared in 12.5% Captisol adjusted to pH 4. ABT-263 and ABT-199 was administered by oral gavage (50 mg/kg). Birinapant was administrated by intraperitoneal injection (15 mg/kg).

#### Bleomycin-induced pulmonary fibrosis model

8-to 10-week-old C57BL/6 male mice were anesthetized and intratracheally instilled with 50 μL of sterile saline (control group) or USP grade bleomycin (1.8U/kg). Body weights were monitored throughout the study. Bleomycin-treated animals with less than 10% weight loss over the first 7 days were excluded from the study. Mice were treated with 50 mg/kg of ABT-199 and 15 mg/kg of birinapant, or vehicle control for 5 days starting 7 days after bleomycin instillation. Animals were euthanized at 14 days after bleomycin injury, and lungs were harvested for qRT-PCR and histological analysis as described below.

#### Non-alcoholic steatohepatitis (NASH) model

6-week-old C57BL/6 male mice were fed with a choline-deficient, L-amino acid defined, high-fat diet (CDA-HFD: A06071302i, Research Diets) for 7 weeks. After 4 weeks of feeding, mice were treated with 50 mg/kg of ABT-199 and 15 mg/kg of birinapant, 50 mg/kg of ABT-263 or vehicle control seven times in 3 weeks. After 7 weeks of high-fat diet, livers were harvested for qRT-PCR analysis as described below, and blood was collected and analyzed by Stanford Animal Diagnostic Laboratory.

### METHOD DETAILS

#### Genome-wide Death-seq positive selection cell death CRISPR screening

For the genome-wide ABT-263 CRISPR–Cas9 screen, the CRISPR/Cas9 deletion library with ten sgRNAs per gene was synthesized, cloned, and infected into Cas9-expressing IMR-90s as previously described (Morgens et al., 2017). ∼500 million IMR-90s stably expressing EF1α-Cas9-BLAST were infected with the genome-wide sgRNA library at an MOI < 1. Infected cells underwent puromycin selection (1.5 μg/ml) for 4 days, after which puromycin was washed out and replaced with normal growth medium. Deep sequencing after selection confirmed sufficient sgRNA library representation. Cells were split into two technical replicates and maintained for ∼2 weeks before inducing senescence by adding doxorubicin at 250 nM for the first 24 hours then 83.3 nM for the subsequent 48 hours before washing out doxorubicin for 4 days in normal growth medium. 7 days after doxorubicin addition, the ∼350 million senescent cells in each technical replicate at ∼1,500x coverage (∼1,500 cells containing each sgRNA) were treated with 1 μM ABT-263 for 24 hours. To harvest DNA from the dying cells, the medium was collected from above the adherent cells to collect detached cells. The tissue culture plates were washed two times in PBS and both washes were also collected and combined with the media. The plates were washed one more time in PBS and then treated with DNase I (Worthington Biochemical Corporation) for 10 minutes at 37°C to remove any remaining genomic DNA from dying cells as described (Kramer et al., 2018) and then washed three more times in PBS to remove residual DNase I and dying cells. The collected media containing dying cells was centrifuged for 5 minutes at 550*g* while the remaining attached living cells on the plates were trypsinized and centrifuged for 5 minutes at 300*g* before genomic DNA was harvested from all screen populations separately by with a DNeasy Blood and Tissue Kit (Qiagen) including proteinase K digestion to inactivate residual DNase I. Deep sequencing on an Illumina Nextseq platform with a NextSeq 500/550 High Output kit was used to monitor library composition which was then directly compared between the dying and live cell populations (“Death-seq”) using casTLE (https://github.com/elifesciences-publications/dmorgens-castle). In brief, the casTLE algorithm compares each set of 10 gene-targeting guides to the non-targeting and “safe-targeting” control sgRNAs. The enrichment of individual guides was calculated as the log ratio between dying and living cell populations, and gene-level effects were calculated from the ten guides targeting each gene. A confidence score (casTLE score) was then derived as a log-likelihood ratio describing the significance of the gene-level effect. P values were then estimated by permutating the targeting guides as previously described (Morgens et al., 2016), and hits were called using FDR thresholds calculated via the Benjamini-Hochberg procedure. See Supplementary Table 1 and Supplementary Table 2 for the complete results for all CRISPR screens. The top 350 genes ranked by casTLE score in the genome-wide ABT-263 CRISPR screen were used to produce a protein-protein interaction enrichment analysis using the Metascape tool (Zhou et al., 2019) and the interaction network was visualized using Cytoscape (Altaf-Ul-Amin et al., 2006). Reactome enrichment analysis was performed by inputting all genes passing a 30% FDR in the ABT-263 genome-wide screen into the Reactome tool (https://reactome.org/PathwayBrowser) (Jassal et al., 2020) and sorting the results on adjusted P value. Reactome IDs for Reactome categories reported in Figure 1 are: R-HSA-111471, R-HSA-109606, R-HSA-111463, R-HSA-111464, R-HSA-111459, R-HSA-111469.

#### Sublibrary Death-seq positive selection cell death CRISPR screening

The targeted custom sublibrary targets a total of 384 genes (10 sgRNA per gene): the top ∼350 genes as ranked by casTLE score from the genome-wide screen as well as select apoptosis and lysosomal-related genes. It also contains 374 control “safe-targeting” sgRNAs. See Supplementary Data 3 for the complete sgRNA composition of the targeted custom sublibrary. Agilent Technologies synthesized the sublibrary oligonucleotides which were cloned into pMCB320 using BstXI/BlpI overhangs after PCR amplification. To conduct head-to-head comparisons of Death-seq versus traditional negative selection screening, both our targeted custom sublibrary and the 10 sgRNA/gene apoptosis and cancer sublibrary were used in screens following the genome-wide screen protocol described above with the exception of the addition of the DMSO-treated control arm, where half the plates of cells in each technical replicate were treated with DMSO instead of ABT-263. The head-to-head screens also had a different screen coverage (in each technical replicate, ∼8,000x coverage for the targeted custom sublibrary head-to-head screens and ∼1,500x coverage for the 35 apoptosis and cancer sublibrary). To conduct parallel screens testing Death-seq with different small molecules and a different inducer of senescence, the genome-wide screen protocol described above was followed except with the targeted custom sublibrary and with a higher coverage of ∼20,000x in each technical replicate. The four treatment groups included one in H_2_O_2_-induced senescent IMR-90s with 24 hours of 1 μM ABT-263 treatment and three in doxorubicin-induced senescent IMR-90s with 24 hours of treatment with 1 μM ABT-263, 10 μM ABT-199, or 2 μM birinapant in each respective treatment arm. Subcellular localizations of proteins encoded by selected screen hits were taken from UniProtKB (The UniProt Consortium, 2021) and MitoCarta3.0 (Rath et al., 2021). MitoCarta3.0’s biological pathway classifications were used to determine the apoptosis pathway members.

#### Lentivirus production and infection

HEK293T cells were transfected with third-generation packaging plasmids and individual sgRNA vectors or sgRNA libraries using PEI max (Polysciences, 26008-5). Lentivirus was harvested after 48 hours and 72 hours and filtered through a 0.45-μm polyethersulfone (PES) filter (Corning 431096). 4 μg/mL of polybrene was used to deliver sgRNA vectors and libraries into IMR-90s. To validate results of genome-wide and sublibrary screens, we infected Cas9-expressing IMR-90 cells with plasmids expressing a single specific sgRNA and puromycin resistance. Two days after infection, the infected cells were puromycin-selected for 4 days and allowed to recover for at least 2 days before experimentation. To additionally validate the PPTC7 screen hit, doxorubicin-induced senescent IMR-90s were infected with plasmids expressing a 10 single specific shRNA and puromycin resistance (see Supplementary Table 4 for sgRNA/shRNA sequences).

#### Cell viability and drug synergy

The cells were plated in triplicate in 96-well plates (Corning, 3603) (typically 7000 senescent and 2000 proliferative cells). Unless otherwise indicated, cell viability was assessed after 3 days of drug treatment, using the XTT assay with the Cell Proliferation Kit II according to the manufacturer’s protocol. Experimental values were normalized using a vehicle-treated and a puromycin-treated (10 μg/ml) condition to set the maximal and minimal viability values, respectively. Drug synergy was assessed as previously described (Han et al., 2017). In brief, drug synergy was assessed using the Bliss independence model (Bliss, 1939). Using the equation F_A_+F_B_ – (F_A_ x F_B_), where F_A_ and F_B_ are the fractional viability inhibitions of the individual drugs A and B at a given dose, the drug combination’s predicted fractional viability inhibition is calculated. The Bliss excess is the difference between the expected viability inhibition and the observed inhibition from the drug combination. Bliss sum is the sum of individual Bliss scores in each matrix of drug doses. Bliss scores greater than zero, close to zero, and less than zero denote synergy, additivity, and antagonism, respectively.

#### Apoptosis assays

To assess apoptosis, Annexin V assays and caspase 3/7 activation assays were used. For the Annexin V assays, APC PI Annexin V solutions were utilized according to the manufacturer’s protocol. In brief, after treatment of the cells with the indicated doses of drugs including some conditions with 20 μM Q-VD-OPH, cells were washed and stained with 1:20 APC Annexin V and 1:10 Propidium Iodide Solution for 15 minutes at room temperature. The fluorescence of the cells was measured on a BD-FACS Aria III flow cytometer. In the case of adherent cells, detached dying cells in the media following drug treatment were centrifuged down and combined with the remaining adherent cells, which were trypsinized. For the caspase 3/7 activation assay, the EarlyTox Caspase-3/7-D NucView 488 assay kit was utilized according to the manufacturer’s protocol. IMR-90s were treated with the indicated doses of vehicle or drugs and 10 μM caspase substrate before reading out fluorescence on a plate reader.

#### Mitochondria membrane potential

Mitochondrial membrane potential was measured using JC-1 staining solution according to the manufacturer’s protocol. IMR-90s were treated with small molecules then stained with JC-1 solution for 5 minutes and the fluorescence of the cells was measured on a BD-FACS Aria III flow cytometer.

#### Quantitative RT-PCR

RNA was extracted from murine lungs and livers using the RNeasy Plus Mini Kit, and cDNA was prepared using High-Capacity cDNA Reverse Transcription Kit according to the manufacturer’s instructions. qPCR was carried out on an ABI 7900HT thermocyler using TaqMan Fast Advanced Master Mix (Applied Biosystems, 4444557) with following Taqman gene expression assays (Applied Biosystems): Il6 (Mm00446190_m1), Hprt (Mm01545399_m1) and Tbp (Mm00446973_m1). The primers and probe used for the detection of *p16*^*INK4a*^ were as follows: p16-F 5′-CGGTCGTACCCCGATTCAG-3′; p16-R 5′- GCACCGTAGTTGAGCAGAAGAG-3′; Probe 5′- [FAM] AACGTTGCCCATCATCA [MGB]-3′.

#### Histopathological evaluation of pulmonary fibrosis

The right lung was inflated and embedded in OCT compound, and 10-μm frozen sections were prepared and stained using hematoxylin and eosin and Masson’s trichrome stain kit according to the manufacturer’s instructions. The severity of pulmonary fibrosis was scored blindly according to the modified Ashcroft scale (Hübner et al., 2008) using Masson’s trichrome stained slides.

### QUANTIFICATION AND STATISTICAL ANALYSIS

Flow cytometry data was analyzed using FlowJo software. Unless otherwise stated, P values were calculated using ANOVA followed by Dunnett’s multiple comparison test to the vehicle group using GraphPad Prism 9 software. Specific data representation details and statistical procedures are also indicated in the figure legends. All data are presented as means ± s.e.m., n.s. = P > 0.05, *P < 0.05, **P < 0.01, ***P < 0.001, ****P < 0.0001.

#### KEY RESOURCES TABLE

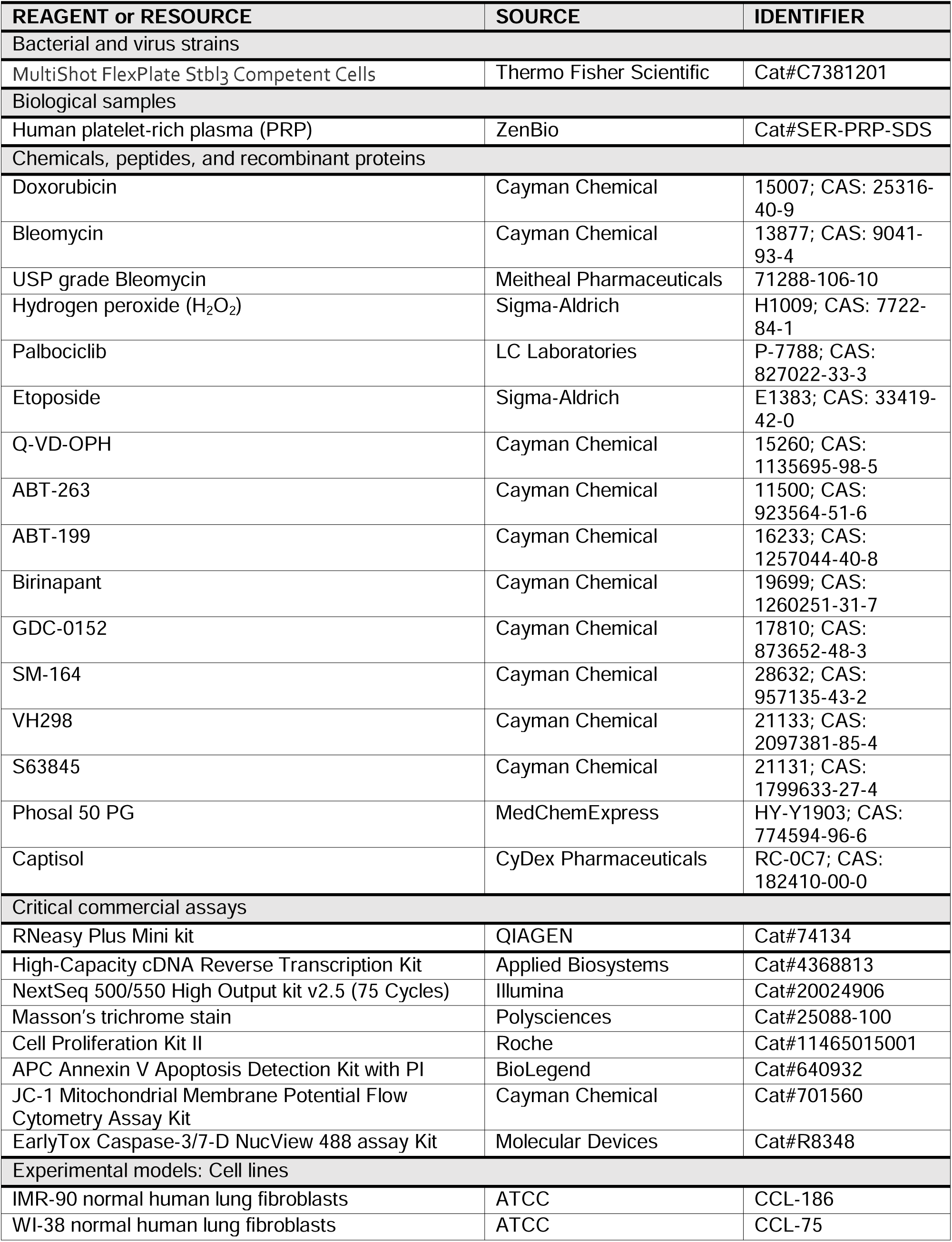
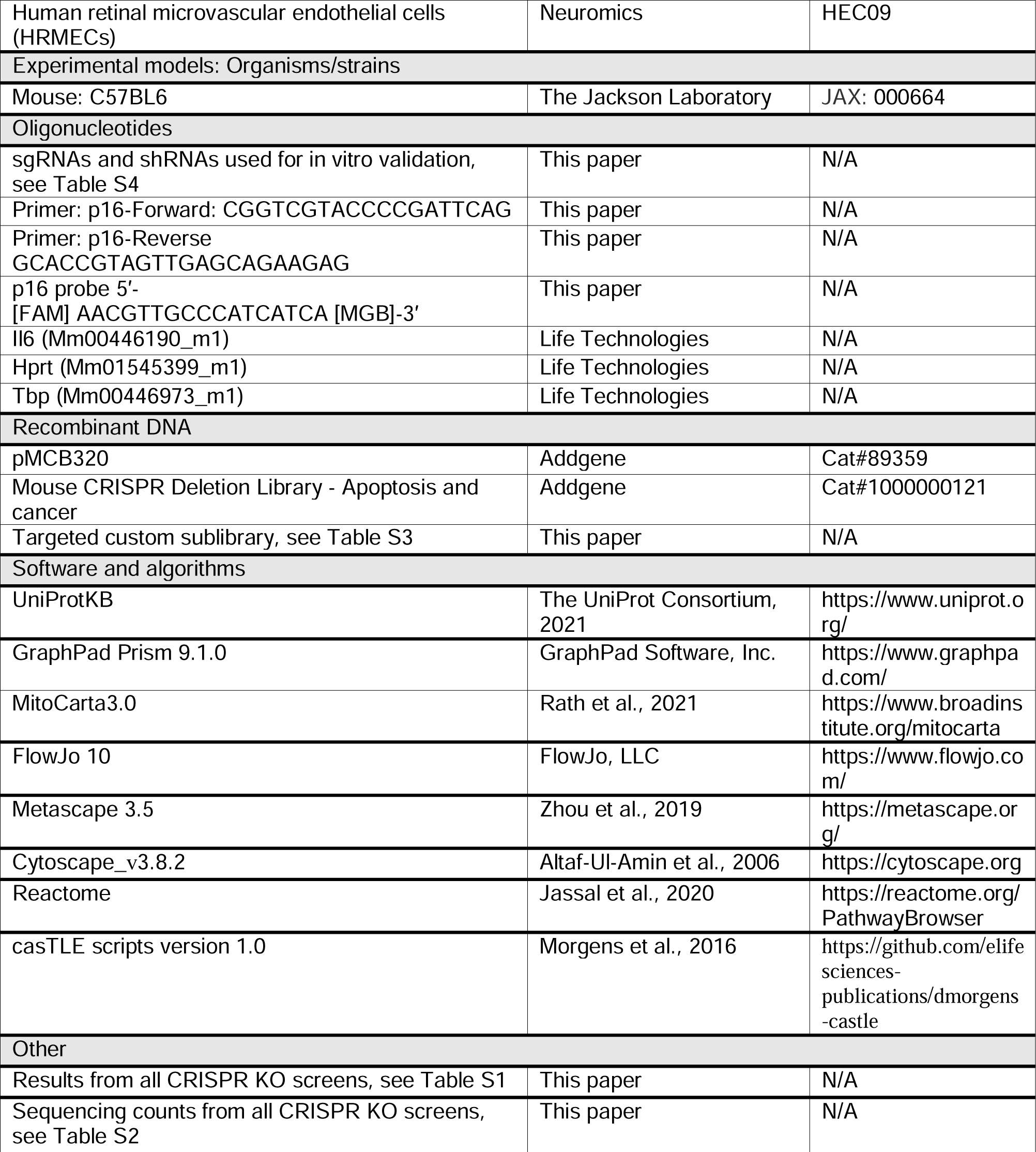

## SUPPLEMENTARY INFORMATION

Supplementary information is available for this paper.

Table S1. Results from all CRISPR KO screens, related to Figures 1, 2, 3, S1, and S2.

Table S2. Sequencing counts from all CRISPR KO screens, related to Figures 1, 2, 3, S1, and S2.

Table S3. Targeted sgRNA composition, related to Figures 2, 3, and S2.

Table S4. Sequences of sgRNAs and shRNAs used in this study.

## SUPPLEMENTAL FIGURE LEGENDS

**Figure S1.**
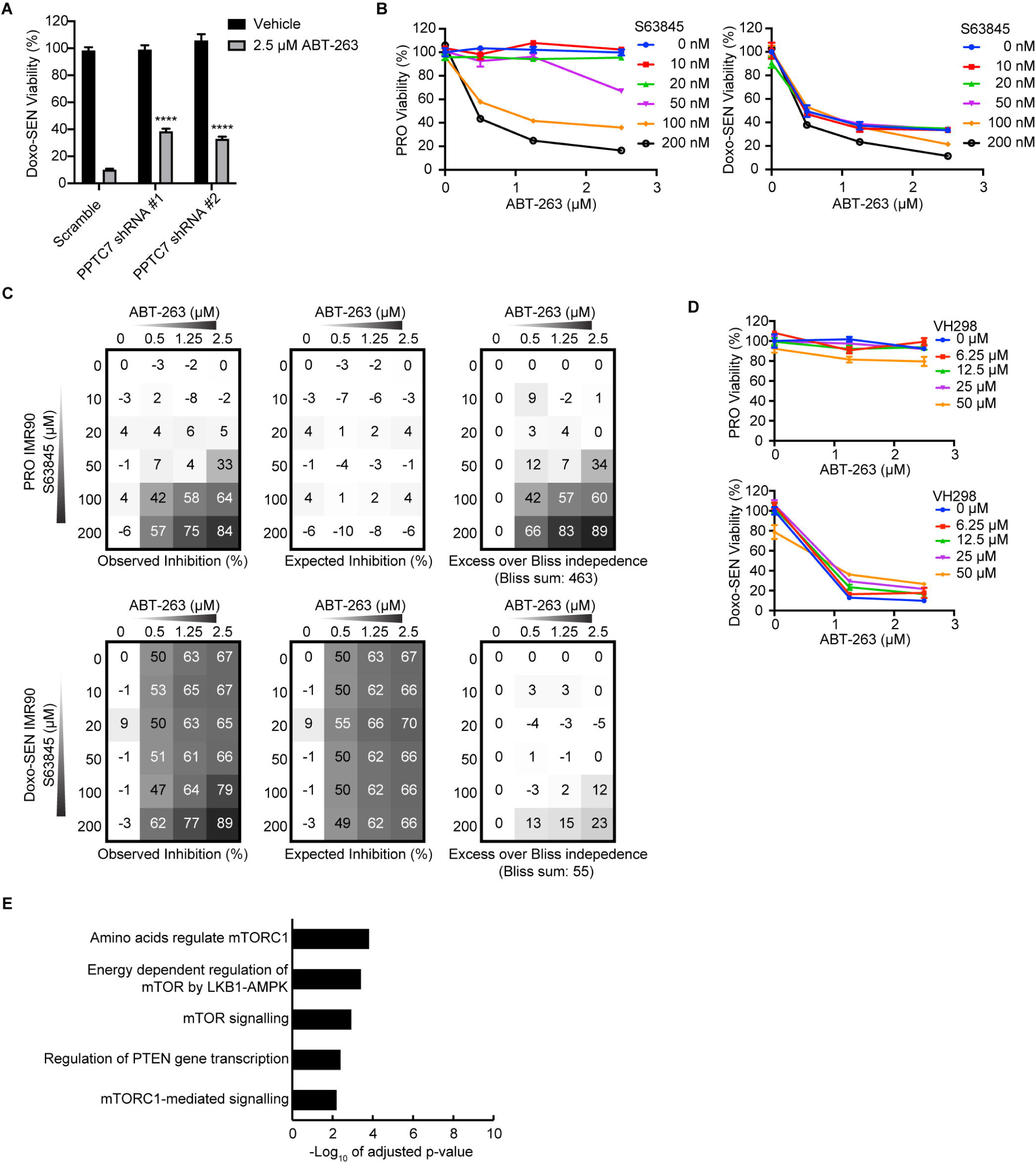
Validations and summaries of Death-seq CRISPR senescence ABT-263 screen, related to Figure 1. (A) Validation of the genome-wide screen ABT-263 resistance modifier hit PPTC7 using individual-well shRNA knockdown compared to control shRNA. Cell viability was assessed after treatment with 2.5 μM ABT-263 for 3 d in Doxo-induced senescent IMR-90s. (B) Proliferative (PRO) (left) and Doxo-induced senescent IMR-90s (right) were treated with ABT-263 and the MCL1 inhibitor S63845 at the indicated concentrations for 3 d before viability was assessed relative to no drug control. (C) The percent expected inhibition is subtracted from the percent observed inhibition at each combination of drug doses in proliferative (top), and Doxo-induced senescent (bottom) IMR-90s to calculate drug synergy represented by excess over Bliss independence. (D) Proliferative (top) and Doxo-induced senescent (bottom) IMR-90s were treated with ABT-263 and the VHL inhibitor VH298 at the indicated concentrations for 3 d before viability was assessed relative to no drug control. (E) The top non-apoptosis related Reactome categories enriched in the 64 genes that passed 30% FDR in the genome-wide ABT-263 screen. Data are representative of two independent experiments performed in triplicate and presented as mean□± □s.e.m. in panels (A), (B), (D). One-way ANOVA with Dunnett’s post-hoc test, *P < 0.05, **P < 0.01, ***P < 0.001, ****P < 0.0001.

**Figure S2.**
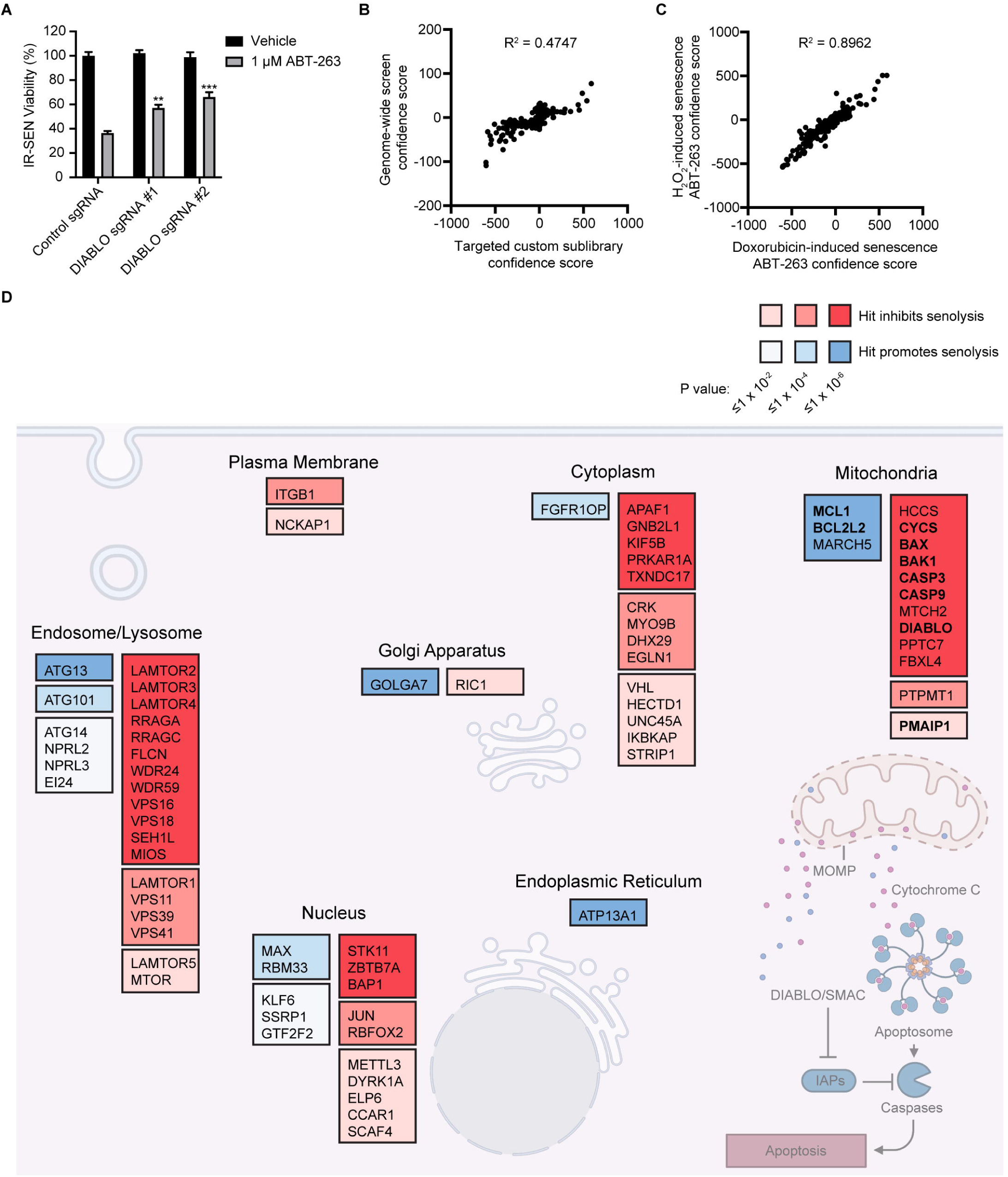
Targeted sublibrary senolytic Death-seq screens highlight importance of SMAC and systematically identify apoptosis pathway members, related to Figure 3. (A) Validation of DIABLO/SMAC sgRNA knockouts compared to control gRNA cell viability after treatment with vehicle or 1 μM ABT-263 for 3 d in IR-induced senescent IMR-90s. (B) Correlation of combo casTLE confidence scores of the targeted custom sublibrary ABT-263 screen and genome-wide screen for all genes in the targeted custom sublibrary. R-squared value is from a linear regression model. (C) Correlation of combo casTLE confidence scores of the targeted custom sublibrary Doxo-induced senescence ABT-263 screen and the targeted custom sublibrary H_2_O_2_-induced senescence ABT-263 screen for all genes in the targeted custom sublibrary. R-squared value is from a linear regression model. Data are representative of two independent experiments performed in triplicate and presented as mean□± □s.e.m. in panel (A). One-way ANOVA with Dunnett’s post-hoc test, *P < 0.05, **P < 0.01, ***P < 0.001, ****P < 0.0001. (D) Schematic of the proteins encoded by selected hits (10% FDR) from the Doxo-SEN custom sublibrary ABT-263 screen with subcellular localizations according to UniProtKB and MitoCarta3.0. In bold are hits that are members of the MitoCarta3.0 apoptosis MitoPathway. Hits are color coded by significance (casTLE P values).

**Figure S3.**
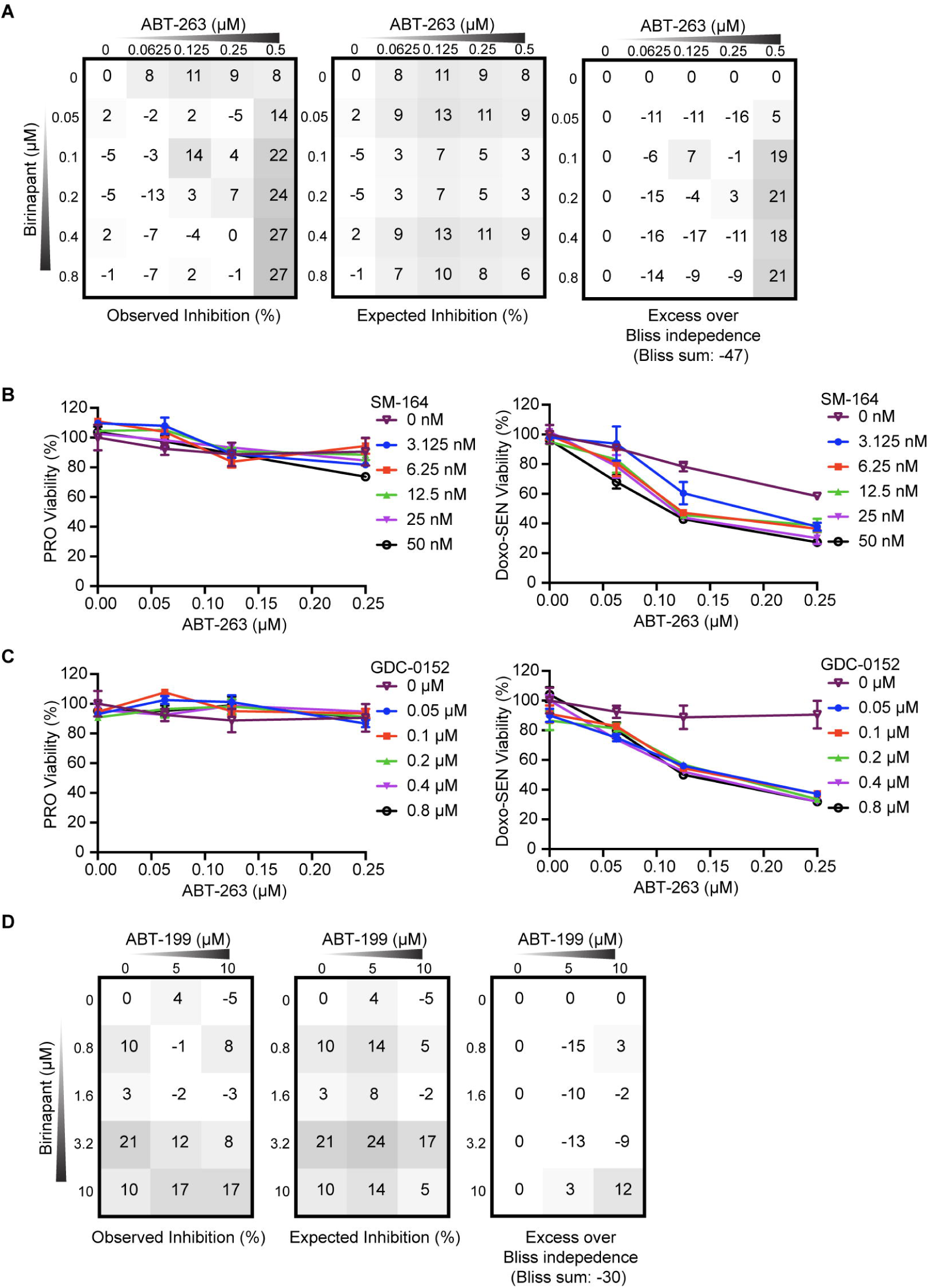
SMAC mimetics synergize with ABT-263 and ABT-199 to induce death in senescent cells, related to Figure 4. (A) The percent expected inhibition is subtracted from the percent observed inhibition at each combination of drug doses in proliferative IMR-90s to calculate drug synergy represented by excess over Bliss independence. (B and C) Proliferative (left) and Doxo-induced senescent (right) IMR-90s were treated with ABT-263 and the indicated SMAC mimetic at the indicated concentrations for 3 d before viability was assessed relative to no drug control. (D) The percent expected inhibition is subtracted from the percent observed inhibition at each combination of drug doses in proliferative IMR-90s to calculate drug synergy represented by excess over Bliss independence. Data are representative of two independent experiments performed in triplicate and presented as mean□± □s.e.m. in panels (A-D).

**Figure S4.**
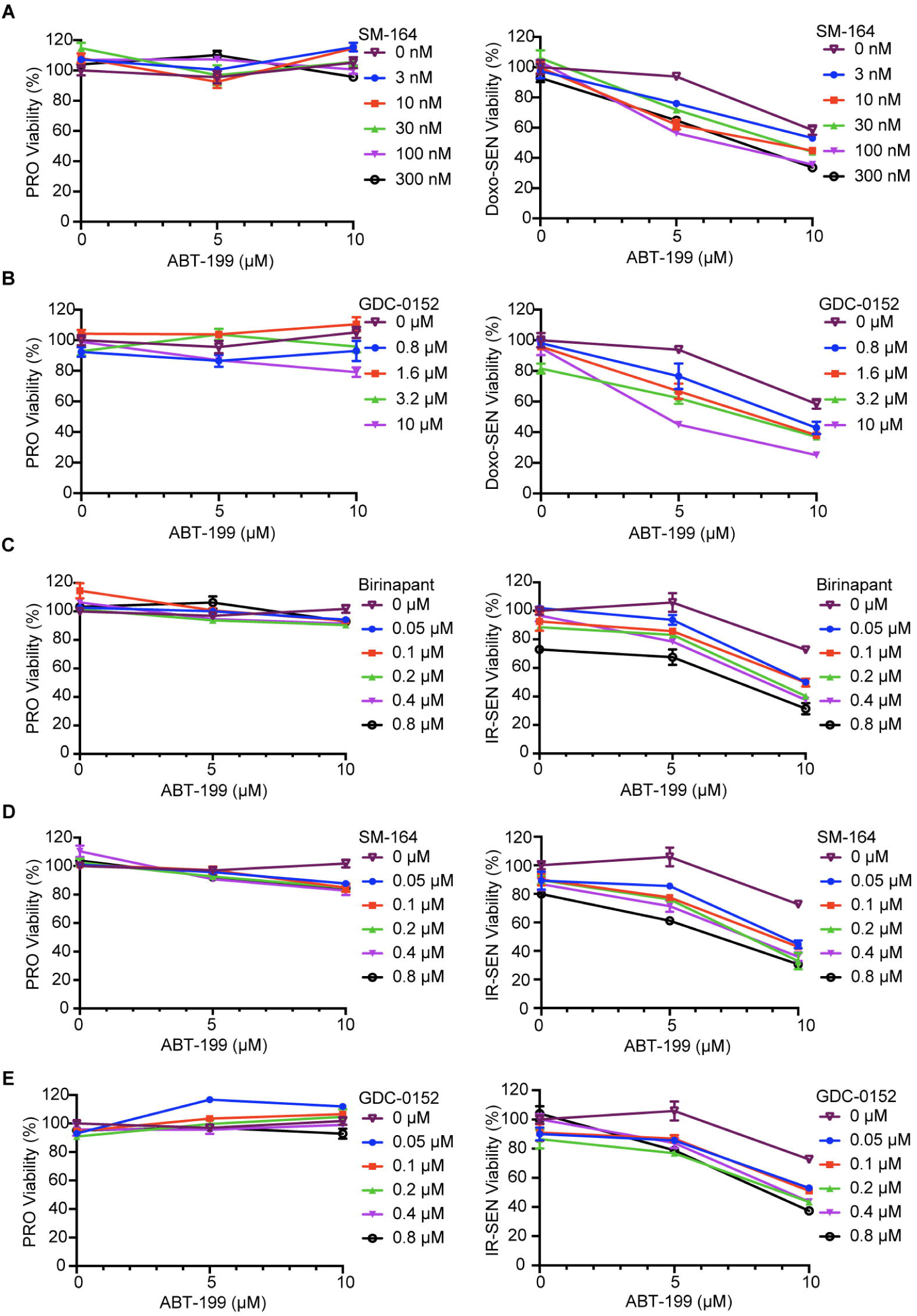
Multiple different SMAC mimetics synergize with ABT-199 to induce death in doxorubicin and irradiation-induced senescent cells, related to Figure 4. (A and B) Proliferative (left) and Doxo-induced senescent (right) IMR-90s were treated with ABT-199 and the indicated SMAC mimetic at the indicated concentrations for 3 d before viability was assessed relative to no drug control. (C-E) Proliferative (left) and IR-induced senescent (right) IMR-90s were treated with ABT-199 and the indicated SMAC mimetic at the indicated concentrations for 3 d before viability was assessed relative to no drug control. Data are representative of two independent experiments performed in triplicate and presented as mean□± □s.e.m. in panels (A-E).

**Figure S5.**
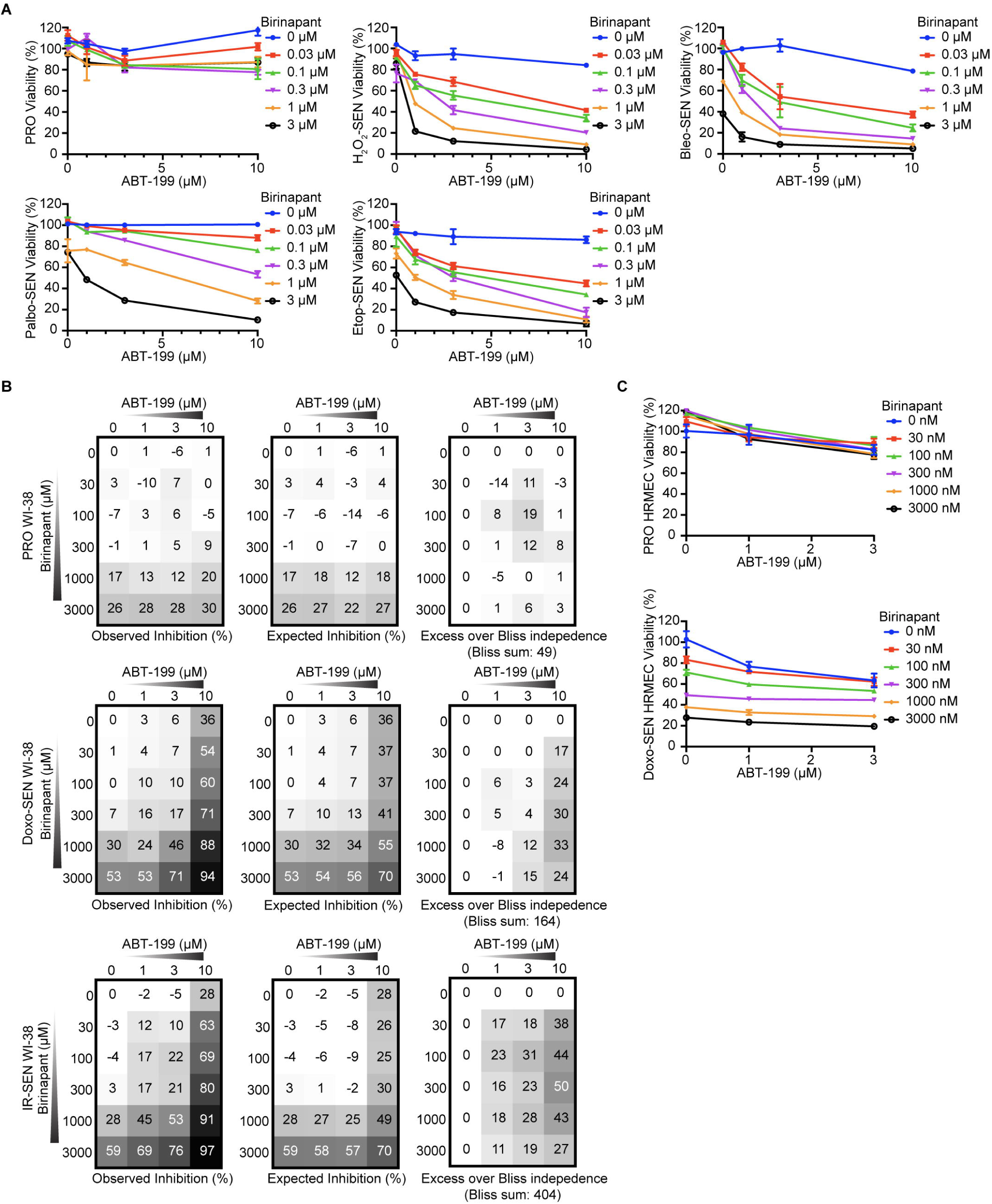
SMAC mimetics synergize with ABT-199 to induce selective senescent cell death across different senescence inducers and cell types, related to Figure 5. (A) Proliferative and senescent IMR-90s induced by treatment with hydrogen peroxide (H_2_O_2_), bleomycin (Bleo), palbociclib (Palbo), and etoposide (Etop) as indicated were treated with ABT-199 and birinapant at the indicated concentrations for 3 d before viability was assessed relative to no drug control. (B) The percent expected inhibition is subtracted from the percent observed inhibition at each combination of drug doses in proliferative (top), Doxo-induced senescent (middle), and IR-induced senescent (bottom) WI-38s to calculate drug synergy represented by excess over Bliss independence. (C) Proliferative (top) and Doxo-induced senescent (bottom) HRMECs were treated with ABT-199 and birinapant at the indicated concentrations for 3 d before viability was assessed relative to no drug control. Data are representative of two independent experiments performed in triplicate and are presented as mean□± □s.e.m. in panels (A-C).

**Figure S6.**
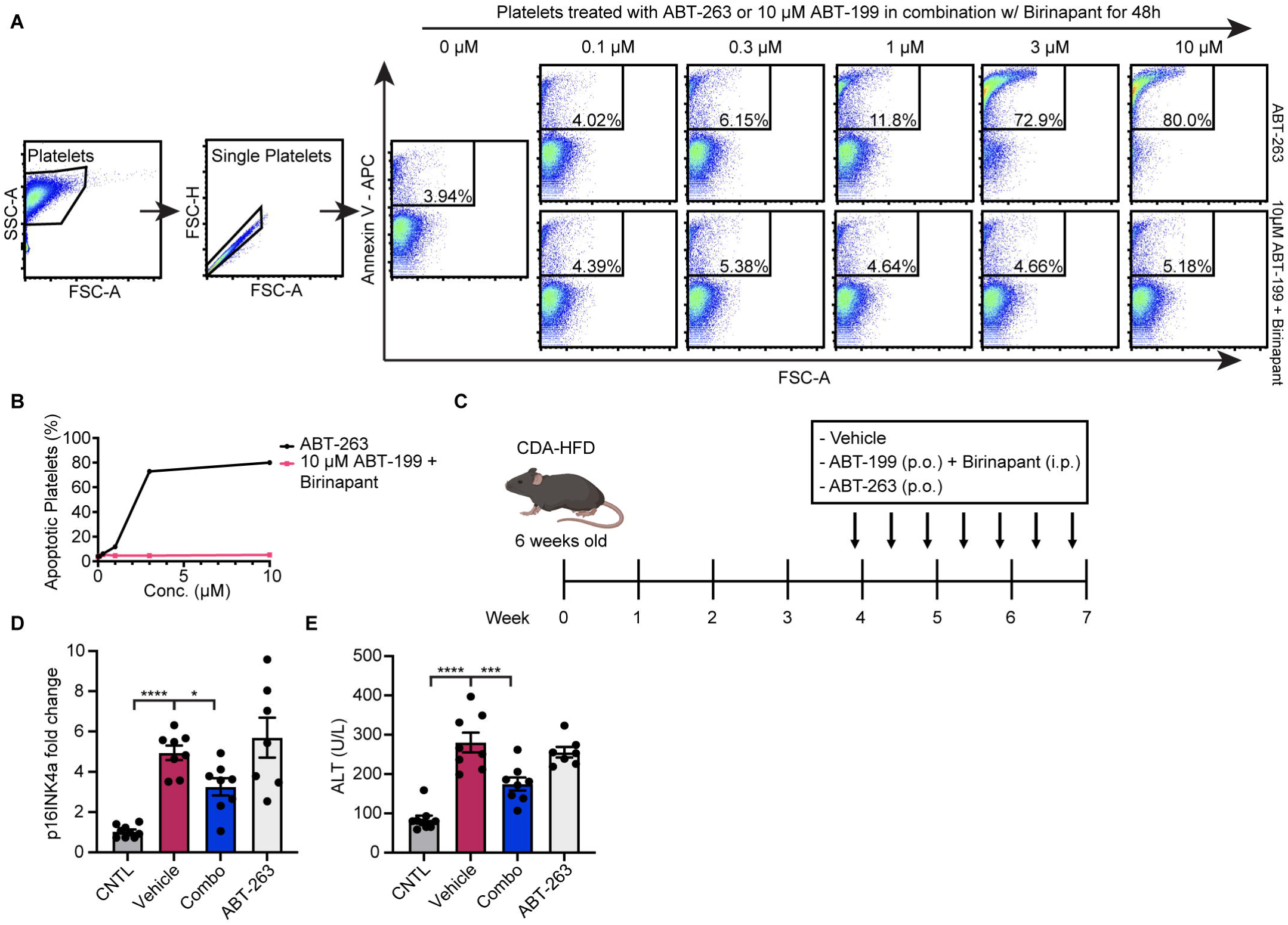
ABT-199 and SMAC mimetic combination spares human platelets and reduces levels of senescent cell markers in vivo in a model of NASH, related to Figure 6. (A) Representative flow cytometry plots of apoptosis in human platelets indicated by Annexin V staining after treatment with vehicle, the indicated doses of ABT-263, or the combination of 10 μM ABT-199 and the indicated doses of birinapant for 48 h. (B) Quantification of (A). (C) Schematic of NASH mouse model experiment. (D) mRNA expression of *p16*^*INK4a*^ in murine livers were quantified by qRT-PCR and normalized by TBP levels. Fold-increase was calculated with respect to the mRNA levels in control mice. (E) Serum alanine aminotransferase (ALT) levels were determined from NASH mice as treated in (C). For panels (D and E) n = 9 control group, n = 8 vehicle and combo groups, and n = 7 ABT-263 group. One-way ANOVA with Dunnett’s post-hoc test, *P < 0.05, **P < 0.01, ***P < 0.001, ****P < 0.0001.

